# PI16 is a non-neuronal regulator of neuropathic pain

**DOI:** 10.1101/696542

**Authors:** Pooja Singhmar, Ronnie The Phong Trinh, Jiacheng Ma, XiaoJiao Huo, Bo Peng, Cobi J Heijnen, Annemieke Kavelaars

## Abstract

Chronic pain is a major clinical problem of which the mechanisms are incompletely understood. Here we describe the concept that PI16, a protein of unknown function mainly produced by fibroblasts, controls neuropathic pain. The spared nerve injury model of neuropathic pain increases PI16 protein levels in fibroblasts in DRG meninges and in the epi/perineurium of the sciatic nerve. We did not detect PI16 expression in neurons or glia in spinal cord, DRG and nerve. Mice deficient in PI16 are protected against neuropathic pain. In vitro, PI16 promotes trans-endothelial leukocyte migration. In vivo, PI16−/− mice show reduced endothelial barrier permeability, lower leukocyte infiltration and reduced activation of the endothelial barrier regulator MLCK and reduced phosphorylation of its substrate MLC2 in response to SNI. In summary, our findings support a model in which PI16 promotes neuropathic pain by mediating a cross talk between fibroblasts and the endothelial barrier leading to barrier opening, cellular influx and increased pain. Its key role in pain and its limited cellular and tissue distribution makes PI16 an attractive target for pain management.

**Graphical Abstract:** 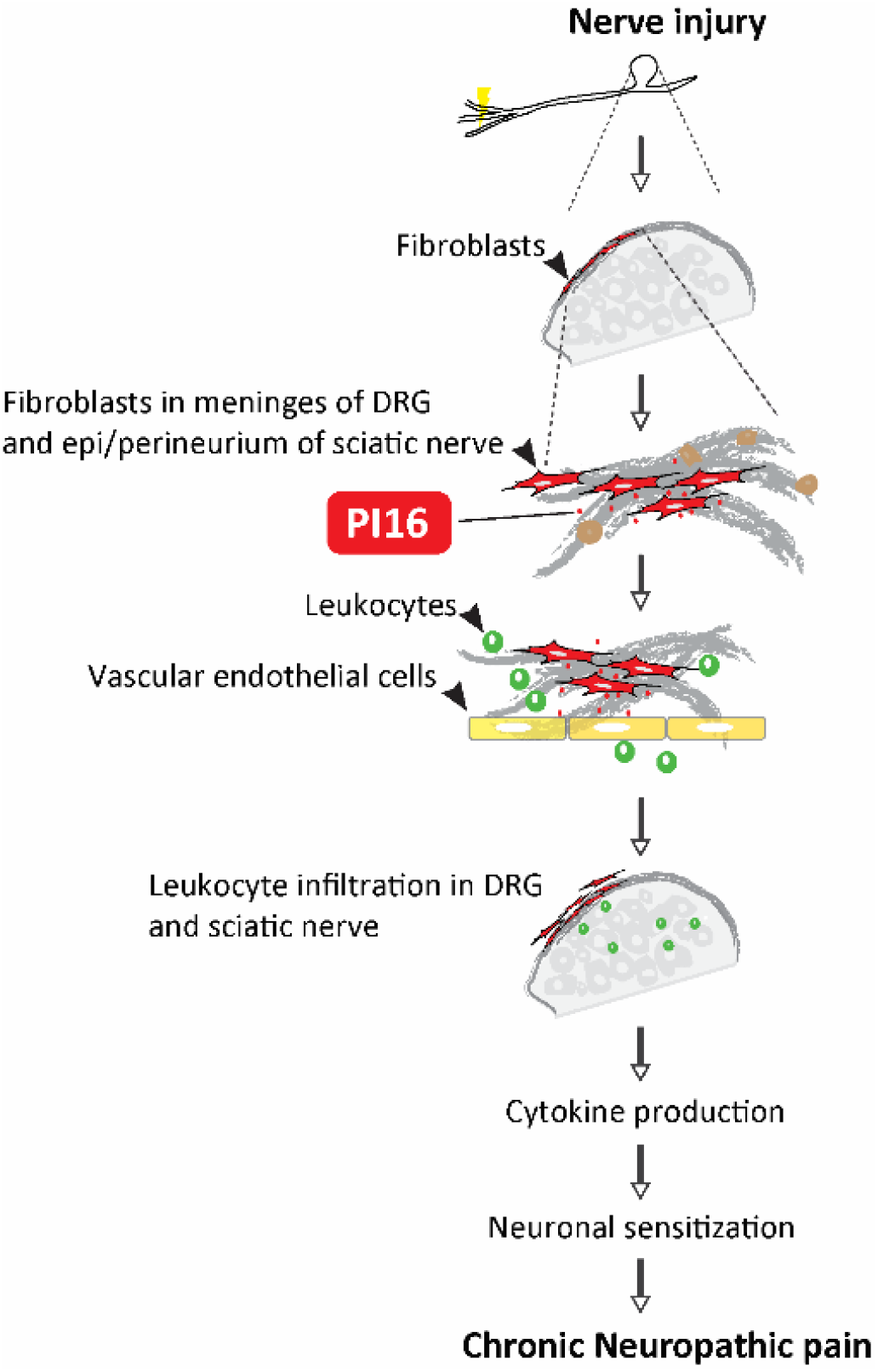

## Introduction

More than one-third of the United States population and approximately 1.5 billion people worldwide face chronic pain^1^. In the United States, the costs of chronic pain are an estimated 30 percent higher than the combined costs of cancer, heart disorders, and diabetes^2^. Treatment of chronic pain is still a major challenge and especially in view of the opioid crisis, better understanding of the cellular and molecular mechanisms underlying chronic pain is urgently needed.

We showed previously that mice deficient in G-protein coupled receptor Kinase 2 (GRK2) develop persistent pain in a number of models^3–6^. In a serendipitous discovery based on RNA sequencing analysis of dorsal root ganglion gene expression comparing GRK-deficient and WT mice, we identified peptidase inhibitor 16 (PI16) as a potential regulator of persistent pain signaling. Pi16 is a poorly characterized member of the CAP (Cysteine-rich secretory proteins, Antigen 5, and Pathogenesis-related 1) superfamily of proteins that is highly conserved across species. PI16 is predicted in the Conserved Domain Database (https://www.ncbi.nlm.nih.gov/cdd) to have an extracellular role and to possess protease enzymatic activity^7^. Very little is known about the tissue distribution and function of PI16. What is known is that PI16 is expressed in cardiac fibroblasts, but PI16-deficient mice do not have a cardiac phenotype^8,9^ There is evidence that Pi16 regulates processing of the chemokine chemerin^9^, cutaneous cathepsin K^10^, and of the matrix metalloprotease MMP2 in endothelial cells^11^. There is no previous evidence for a role of PI16 in pain signaling. Here, we demonstrate for the first time that PI16 plays a key role in chronic pain is expressed in fibroblasts in the meninges of the DRG and the perineurium, but not by neurons or glia.

## Results

### Identification of Pi16 as a novel gene associated with pain

To identify novel pathways that regulate chronic pain, we compared gene expression profiles between DRG from wild type (WT) and GRK2+/− mice at 5 h after intraplantar PGE_2_. We selected this model, because WT mice recover within hours from PGE_2_-induced allodynia, whereas GRK2+/− mice transition to chronic pain and show allodynia for at least 3 weeks^3–5^.

Only 5 differentially expressed genes (Pi16, Gm12250, ligp1, Gbp2, Irgm2) were identified in the DRG (Figure 1A and Figure S1A) after elimination of genes that differed by genotype in the absence of PGE_2_ (Figure S2). Out of these 5 genes we further explored Pi16. We selected Pi16 because it is a gene with unknown function in pain, is highly conserved in eukaryotes and its protein sequence shares high degree of identity with human PI16. The results in Figure 1B demonstrate that Pi16 mRNA expression was decreased in DRG from WT mice during recovery from PGE_2_ allodynia, but did not change in DRG from GRK2+/− mice who developed persistent allodynia for at least 21 days after PGE_2_ (Figure 1B). In a separate group of WT mice we validated our findings and showed that Pi16 mRNA levels decrease during resolution of PGE_2_-induced allodynia and return to base line levels at 48hrs after PGE2 injection (Figure S1B).

**Figure 1.**
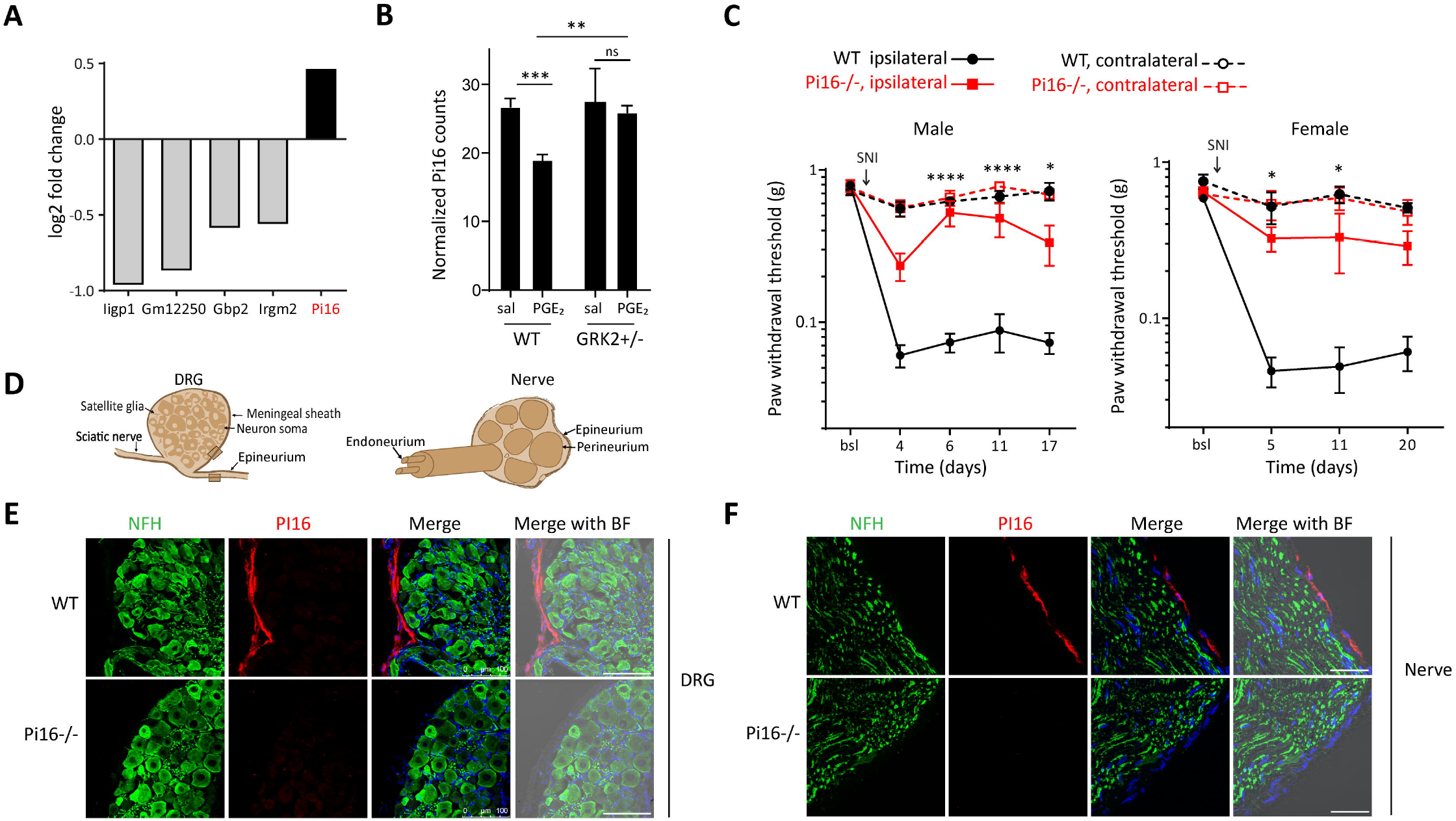
Identification of Pi16 as a novel gene associated with pain and PI16 is localized in the meninges of DRG and the epi/perineurium. **(A)** Differentially expressed genes as identified by RNA-sequencing analysis of lumbar DRGs from WT (n=3) and GRK2+/− (n=3) mice collected 5 hours after PGE_2_ treatment (100 ng/paw). Fold change is based on counts per million reads adjusted for multiple testing error (False discovery rate < 0.05) and a positive fold change indicates higher levels in GRK2+/− mice treated with PGE_2_ (also see Figure S1). Genes that were differentially expressed when comparing saline-treated WT and GRK2+/− mice were eliminated from the analyses. **(B)** Expression of Pi16 mRNA in lumbar DRGs of saline (sal) and PGE_2_ treated WT (n=3) and GRK2+/− (n=3) mice 5 hours after PGE_2_ treatment as assessed by RNA-seq analysis. Data represent normalized log-counts-per-million reads. ***P < 0.001, **P < 0.01; two-way ANOVA test, post hoc Bonferroni analysis. **(C)** Spared nerve injury (SNI) was performed on (C) male WT (n=6) and Pi16−/− (n=4), and, **(D)** female WT (n=4) and Pi16−/− (n=5) mice. Changes in 50% paw withdrawal threshold (mechanical allodynia) were measured using von Frey hairs in ipsilateral (left) and contralateral hind paw at baseline (bsl) and over time after SNI. Two-way repeated-measures ANOVA: (C) main effect of time (p< 0.0001), genotype (p< 0.001), and a genotype-by-time interaction (p< 0.001). post hoc Bonferroni analysis: *p<0.05, **** p<0.0001 Pi16−/− ipsilateral vs WT ipsilateral; (D) main effect of time (p< 0.0001), genotype (p< 0.001), and a genotype-by-time interaction (p< 0.045). post hoc Bonferroni analysis: *p<0.05, Pi16−/− ipsilateral vs WT ipsilateral. (D) Schematic illustration of DRG and sciatic nerve containing neuronal soma, glia, blood vessels. The DRG is enclosed by a meningeal sheath that is continuous with the epineurium of the sciatic nerve. The epineurium surrounds the entire nerve and is present between the nerve fascicles. The two other connective tissue compartments in the nerve are the perineurium (around each fascicle) and endoneurium (around the axons). Shaded box is to orient to the region where the images in panel B and C were captured. **(E)** Representative lumbar DRG section from naïve WT and Pi16−/− mice stained with neuronal marker, NFH and PI16. PI16 immunostaining is detected only in the meningeal sheath around the DRG and not in the neuronal soma or in other cells inside the DRG. Scale bar indicates 100 μm. **(F)** Representative sciatic nerve sections from WT and Pi16−/− mice showing immunostaining with neuronal marker NFH and PI16. Right most panel in (E) and (F): Bright field (BF) image merged with green (NFH), red (PI16) and DAPI (blue) channel is shown. Scale bar indicates 100 μm.

### Pi16 knock out mice are protected against SNI-induced pain

To determine the role of Pi16 in chronic pain, we used the spared nerve injury (SNI) model of chronic neuropathic pain in Pi16−/− and WT mice. Male and female Pi16−/− mice were protected against SNI-induced mechanical allodynia (Figure 1C). Mechanical sensitivity at baseline and in the contralateral paw after SNI was not affected by genetic deletion of the Pi16 gene.

Protection of Pi16−/− mice against neuropathic pain was not due to changes in the distribution of subtypes of DRG neurons or spinal cord dorsal horn neuron staining for NF200, CGRP or IB4 (Figure S3). The morphology and structure of myelinated and unmyelinated fibers in the sciatic nerve and the density of PGP+ intra-epidermal nerve fibers also did not differ between WT and PI16 KO mice (Figure S3). These findings indicate that genetic deletion of Pi16 does not result in structural abnormalities in the peripheral nervous system. Together these findings led us to conclude that Pi16 plays a key role in neuropathic pain.

### PI16 is detected in fibroblasts in the meninges of the DRG and the epi/perineurium

We next characterized localization of PI16 protein in DRG and sciatic nerve. Interestingly, under base line conditions, PI16 protein is expressed in the meningeal sheath surrounding the DRG but is not detectable in DRG neurons or satellite glia (Figure 1D and E). The meninges of the DRG are continuous with the perineurium which surrounds the nerve fascicles and endoneurial space^12,13^ (Figure 1D). PI16 is clearly detectable in the epi/perineurium of the sciatic nerve, but not in neurons and glia (Figure 1F).

In the meninges and epi/perineurium, PI16 is present in elongated cells with flat and wavy nuclei which is a characteristic of fibroblasts, a major cell type present in dura and arachnoid sheaths. Because of the high degree of heterogeneity among fibroblasts and the lack of a pan-fibroblast marker, we used two different proteins to label fibroblasts; α-SMA (alpha smooth muscle actin) and P4HB (collagen Prolyl 4-hydroxylase). Double-staining for PI16 and a-SMA or P4HB confirmed expression of PI16 exclusively in fibroblasts (Figure 2A). We did not detect any PI16 staining in GLUT-1-positive cells in the inner layer of the perineurium (Figure 2B). There is some evidence that PI16 is expressed in endothelial cells^11^, but we did not detect PI16 in CD31-or CLDN1-positive endothelial cells in the DRG (Figure 2C). However, staining of DRG showed that PI16 positive fibroblasts were closely associated with endothelial cells (Figure 2C, top panel). A whole mount DRG staining to visualize spatial distribution of PI16 protein showed a number of PI16-positive fibroblasts around the CD31 vascular network (Figure 2C, middle panel). PI16 is also expressed in fibroblasts in the meninges of spinal cord (Figure S4). We did not detect PI16 in neurons or glia in the spinal cord itself and PI16 was not detected in fibroblasts surrounding the central canal (Figure S4A-C).

**Figure 2.**
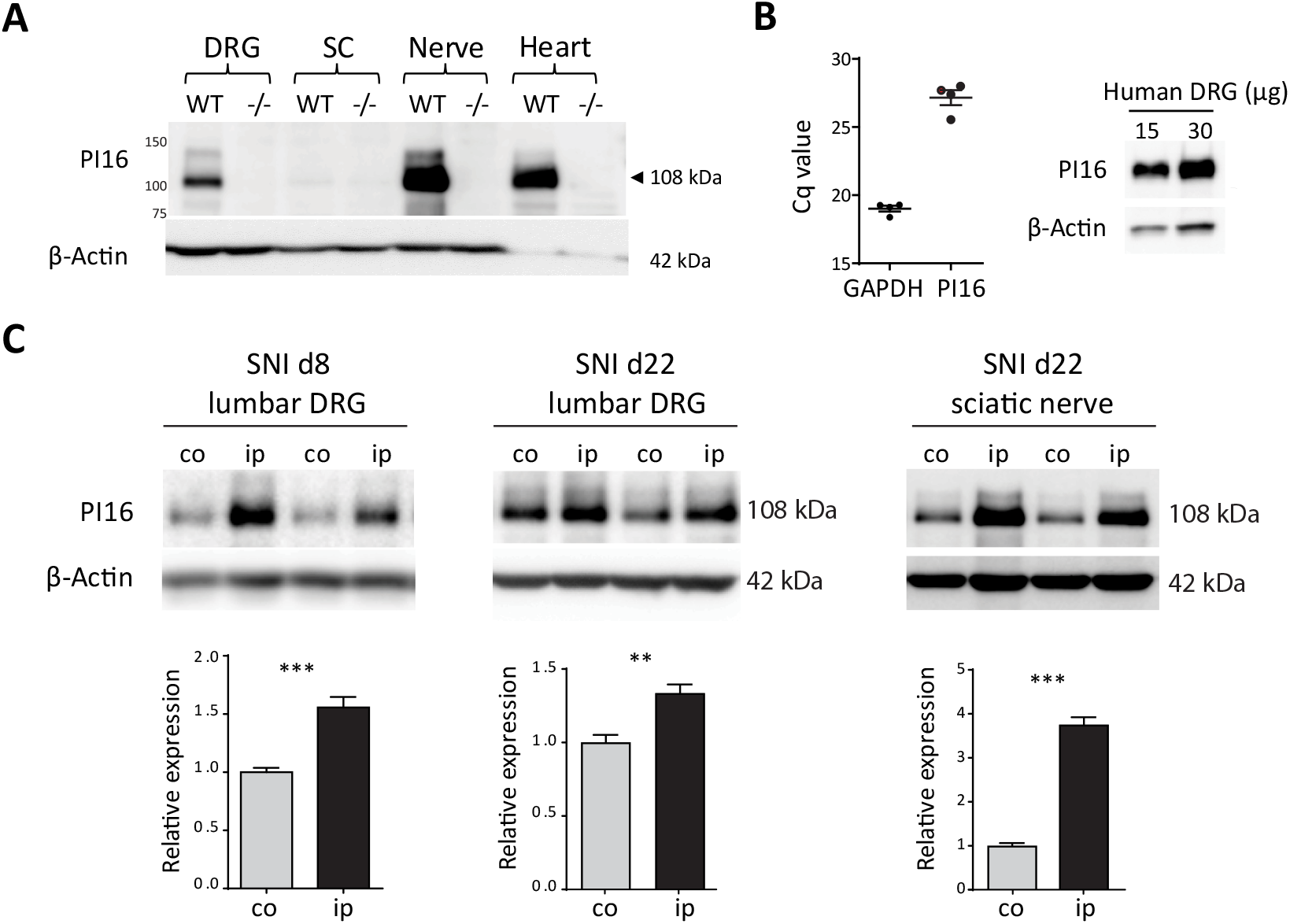
PI16 is detected in fibroblasts in the meninges of the DRG and the epi/perineurium. Representative sciatic nerve section from naïve WT mice showing double immunostaining of PI16 (red) with markers of different cell types (green) present in and around meningeal sheath. **(A)** Fibroblast markers α-SMA and P4HB **(B)** perineurial marker GLUT-1, **(C)** endothelial markers CD31 and CLDN1. Top panel of (C) shows PI16-positive fibroblasts and endothelial cells are closely positioned within 1 μm distance. Middle panel of (C) shows staining of whole mount DRG. Bright field (BF) image merged with other channels is shown. Scale bar indicates 25 μm.

### PI16 protein levels in DRG and sciatic nerve are increased in models of neuropathic pain

PI16 is mainly expressed as a 108 kDa protein in murine DRG, nerve (Figure 3), and in meninges of brain and spinal cord (Figure S4D). No expression of the 108 kDa protein was detected in samples from PI16−/− mice, confirming antibody specificity. The results in Figure 3B show that PI16 mRNA and protein is also expressed in human DRG samples obtained during spinal surgery wherein a spinal nerve root was sacrificed as the standard of care. These findings confirm that expression of PI16 in DRG is conserved between mouse and human.

**Figure 3.**
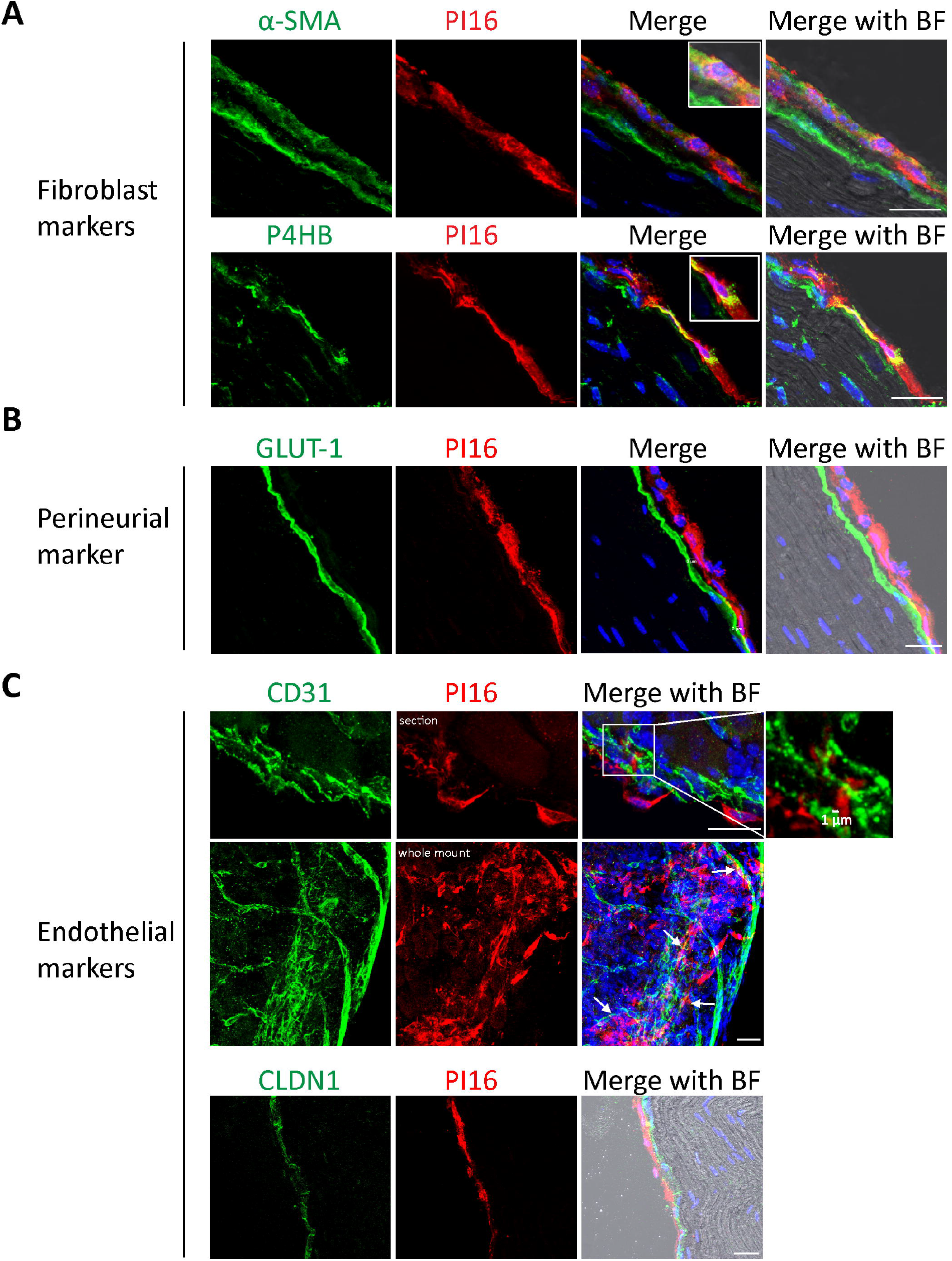
SNI upregulates PI16 protein levels in DRG and sciatic nerve. **(A)** Western blot analysis showing PI16 expression in murine DRG, spinal cord, nerve, and heart. PI16 is mainly expressed as a 108 kDa protein and a faint higher molecular weight band around 140 kDa. Absence of 108 kDa band in samples from Pi16−/− mice confirms the antibody specificity. Note-β-actin antibody used does not react with heart tissue. **(B)** Left panel-Quantitative RT-PCR analysis of PI16 mRNA in human DRG samples. Cq value are plotted and each data point represents one sample. GAPDH was used for normalization. Right panel-Western blot analysis of PI16 protein in a human DRG. Two lanes have different amount of protein from one sample. **(C)** Western blot analysis of PI16 protein expression (detected as a 108kDa band) in ipsilateral (ip) and contralateral (co) lumbar DRG at 8 and 22 days after SNI, and sciatic nerve 22 days after SNI. Representative blots showing PI16 expression in two animals per group. Bar graphs represent means ± SEM of n= 6 mice per group from three independent experiments. **p<0.01, *** p<0.001, (t test).

SNI significantly increased PI16 protein levels in the ipsilateral lumbar DRGs, and this increase was detected both at day 8 and day 22 after SNI (Figure 3C). SNI also induced a marked increase in PI16 protein in the ipsilateral sciatic nerve (Figure 3C). Sham-surgery did not affect PI16 levels (Figure S4E).

Next, we determined in which cell type in DRG meninges and in the epi/perineurium PI16 is increased in response to SNI. SNI markedly increased PI16 staining in the meningeal sheath of the ipsilateral DRG and epi/perineurium of the nerve (Figure 4A and B). However, SNI did not induce detectable PI16 in neurons or glia in the DRG or nerve (Figure 4). The SNI-induced increase in PI16 and its cellular distribution were maintained at all time points tested (day 5, day 22, and day 45 post SNI). SNI did not induce detectable changes in PI16 staining in the meninges of the spinal cord, in the dorsal horn of the spinal cord (Figure S4F and G).

**Figure 4.**
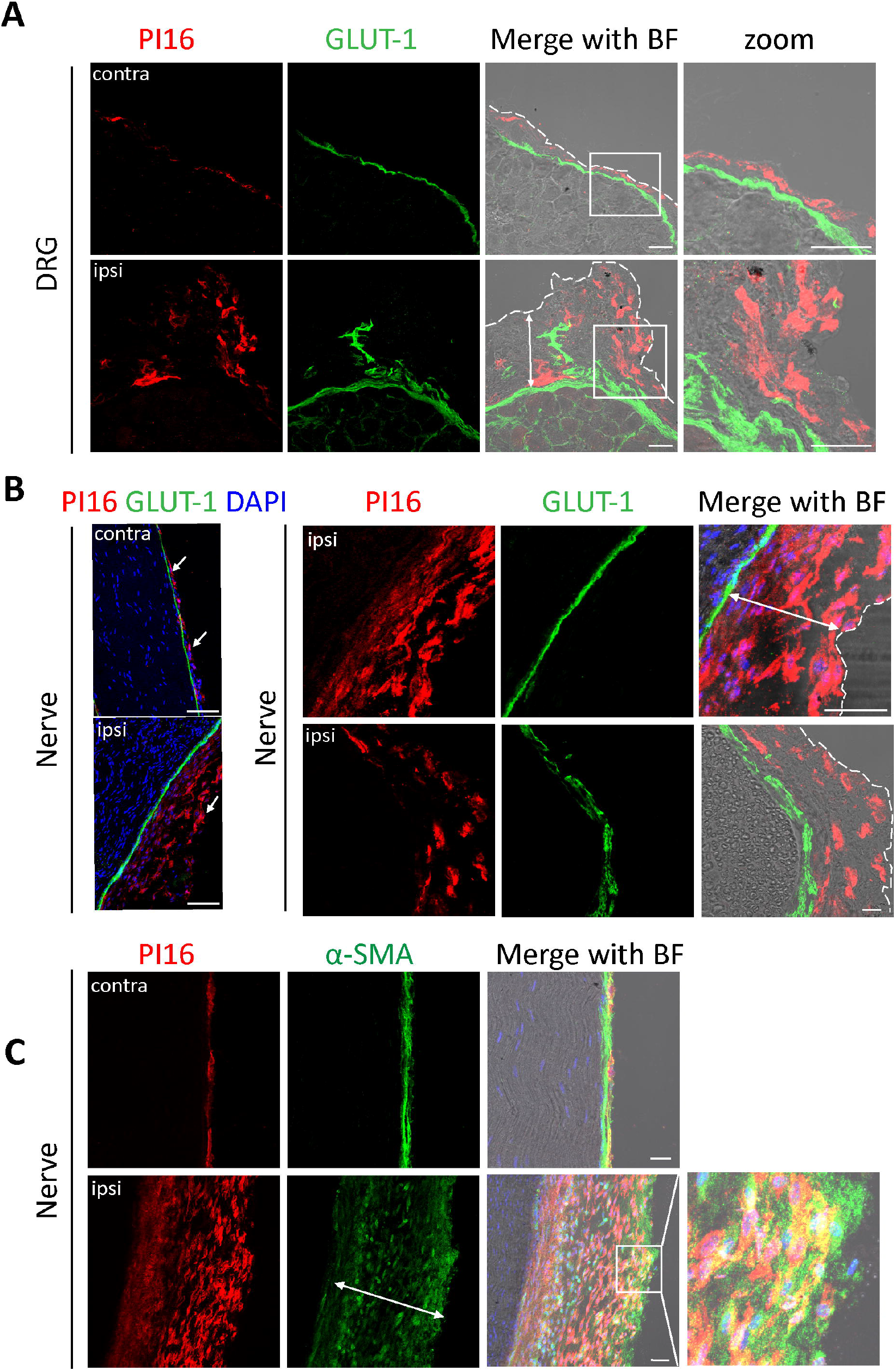
SNI-induced increase in fibroblast Pi16 levels in WT mice. **(A)** Representative image showing immunostaining of PI16 (red) and GLUT-1 (green) in lumbar DRG section from side contralateral (contra) and ipsilateral (ipsi) to SNI surgery. BF (bright field) image merged with PI16 and GLUT-1 is shown. The zoom panel show a magnified view of the area outlined by the square. Perineurial GLUT-1 (green) staining marks the border of DRG and the dotted white line marks the edge of the meningeal sheath. Note the PI16 staining in the meningeal sheath. Note increased PI16 staining outside GLUT-1 in the meninges of ipsilateral DRGs compared to contralateral. Scale bar indicates 25 μm. **(B)** Representative merged images showing immunostaining of PI16 (red), GLUT-1 (green), and DAPI (blue) in sciatic nerve longitudinal section from ipsilateral and contralateral side after SNI. Note increase PI16 staining (arrow) outside GLUT-1 in ipsilateral nerve compared to contralateral. Scale bar indicates 100 μm. Note increased thickness of epi/perineurium on the right panel (double headed arrow) in sciatic nerve longitudinal (top) and cross section (bottom). Scale bar 100 μm (top), 25 μm (bottom). **(C)** Representative image showing PI16 (red) immunostaining in α-SMA (green) positive fibroblast in sciatic nerve from contralateral and ipsilateral side. Note expansion of α-SMA positive fibroblasts co-expressing PI16 in the ipsilateral nerve. BF (bright field) image merged with PI16, α-SMA, and DAPI (blue) is shown. Scale bar indicates 25 μm.

In the DRG and nerve, the SNI-induced increase in PI16 staining was associated with increased meningeal and epi/perineurial thickness and expansion of fibroblasts which were predominantly positive for α-SMA and co-expressed PI16 (Figure 4C). An expansion of PI16 positive fibroblasts was also observed around the injury site in the nerve (Figure 5A). After SNI, different cell types, including immune cells infiltrate into the DRG and nerve. Expression of PI16 by a subset of memory T regulatory cells has been reported^14^. Therefore, we investigated whether the infiltrating leukocytes expressed PI16. However, we did not detect any co-labeling of Pi16 with the pan leukocyte marker CD45 in DRG or at the site of nerve injury, indicating that leukocytes do not contribute to the increase in Pi16 expression in response to SNI (Figure 5B). We also checked for the presence of PI16 in cells positive for chondroitin sulfate proteoglycan (NG2) which labels endoneurial-fibroblasts, pericytes, and vasculature associated muscle cells^15,16^. NG2 positive fibroblasts were detected in the endoneurial space and around the injury site, but these cells did not contain PI16 (Figure 5C).

**Figure 5.**
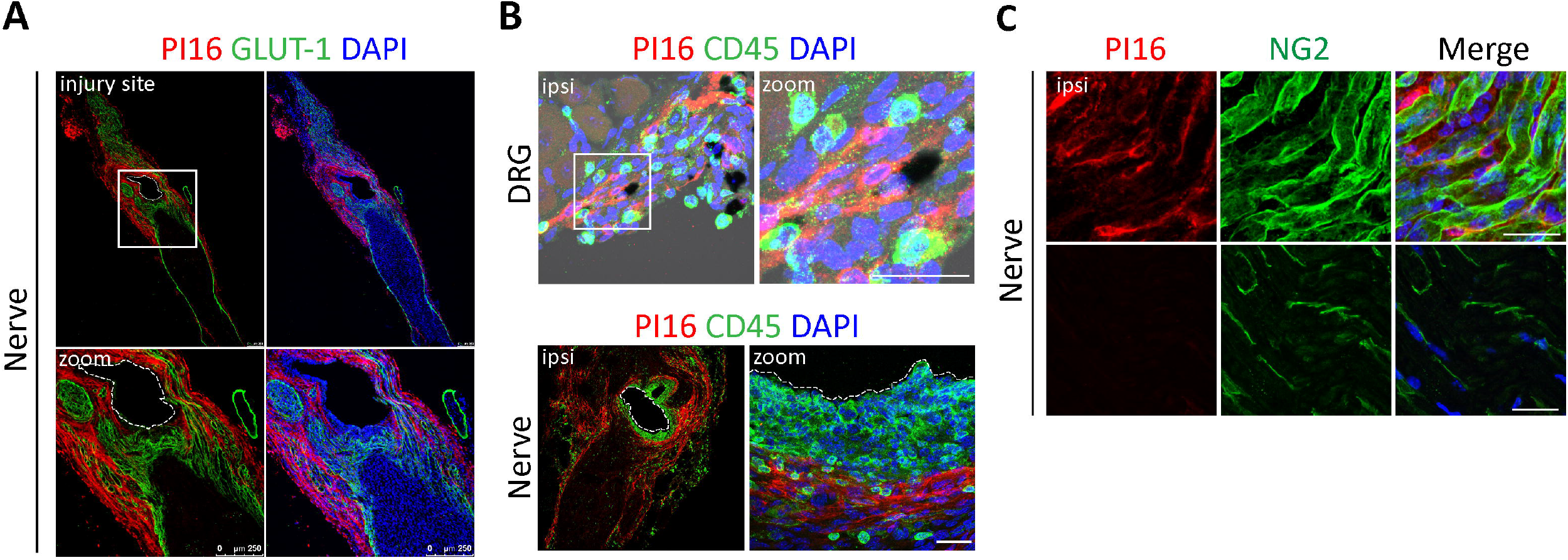
PI16 is not detectable in other cells than fibroblasts. **(A)** Representative merged image showing PI16 (red), GLUT-1 (green) and DAPI (blue) immunostaining around the nerve injury site in the sciatic nerve (day 45 post SNI). White box indicates the injury site from SNI surgery and the bottom panel shows its magnified view. Scale bar indicates 250 μm. **(B)** Representative lumbar DRG and sciatic nerve section from side ipsilateral to SNI surgery showing immunostaining of PI16 (red), CD45 (green), and DAPI (blue) staining. Zoom panel shows a magnified view of the area outlined by the white square (top) and a magnified view of the nerve injury site in the sciatic nerve (bottom). Scale bar indicates 100 μm. **(C)** Representative sciatic nerve section from the injury site showing the endoneurial cell marker NG2 (green), PI16 (red), and DAPI (blue) staining. Top panel shows an area next to the injury site and bottom panel shows endoneurial space. Scale bar indicates 100 μm.

### Pi16 is secreted in vitro by myofibroblasts

The SNI-induced increase in PI16 in the DRG and nerve was mainly detected in α-SMA positive fibroblasts. In vitro, differentiation of primary perineurial fibroblast into α-SMA positive myofibroblasts using TGFβ1 increased Pi16 protein levels in these cells both intracellularly and in the culture supernatant (Figure 6A and B) indicating PI16 is a secreted protein. To assess the subcellular distribution of PI16, we overexpressed GFP-tagged PI16 in fibroblasts (L cells). As expected for a secretory protein, GFP-tagged Pi16 localized to the endoplasmic reticulum and secretory vesicles (Figure 6C).

**Figure 6.**
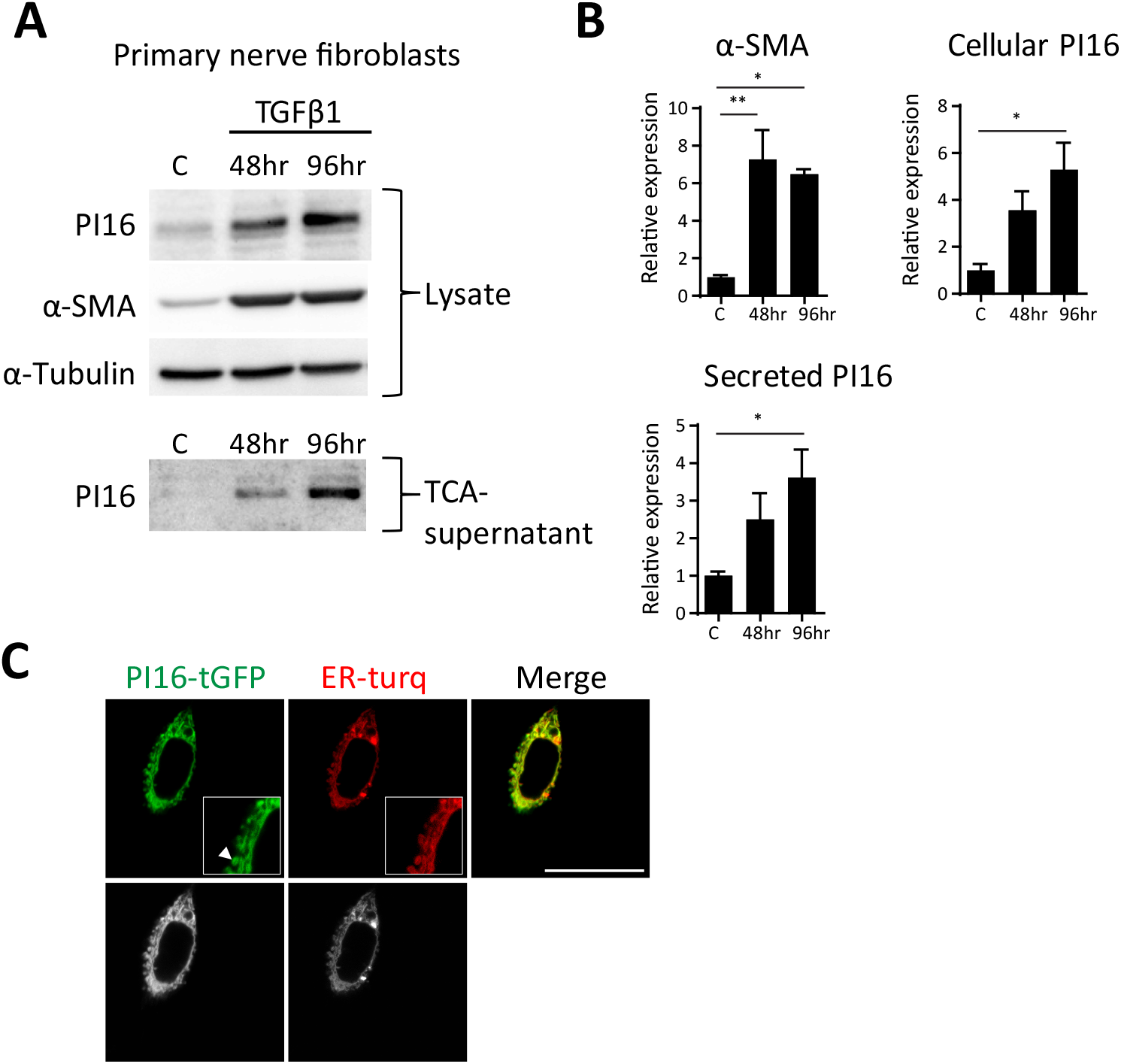
Pi16 is secreted by myofibroblasts in vitro. **(A)** Mouse fibroblasts cultured from postnatal day 8 mouse sciatic nerve were serum starved for 24 hr and treated with TGF-β1 (5ng/ml) for 48 hr or 96 hr followed by Western blot analysis of total cell lysate or TCA precipitated culture supernatant. Representative western blots are shown. α-SMA was used as a myofibroblast marker and α-Tubulin was loading control. **(B)** Quantification of PI16 and α-SMA for panel B. Bar graph depicts means ± SEM of at least three independent experiments. *P < 0.05; **P < 0.01 (analyzed using t test). **(C)** Live imaging of mouse fibroblasts (L cells) transfected with mTurquoise2 (endoplasmic reticulum marker, ER-turq) and PI16 tagged with turbo-GFP. Raw black and white images are shown in bottom panel. Note PI16 staining in endoplasmic reticulum vesicles and tubules (arrowhead) in the magnified view in the white box. Scale bar indicates 25 μm.

### Pi16 deficiency reduces the SNI-induced infiltration of leukocyte into the DRG and nerve

To get insight into the pathways via which Pi16-deficency protects against SNI-induced pain, we compared SNI-induced changes in the transcriptome of lumbar DRGs from WT and Pi16−/− mice by RNA sequencing. In WT mice, SNI changed the expression of 4074 genes in the DRG while only 1138 genes changed in Pi16−/− mice (−2.00<log_2_ Fold Change<2.00), as compared to their sham-surgery control groups. Functional enrichment analysis using Ingenuity Pathway Analysis (IPA) identified activation of canonical pathways associated with neurotransmission and inflammation upon SNI surgery (Figure S5). Notable pathways included neuroinflammation signaling, cAMP signaling, and complement system, which is in agreement with previously published transcriptome data from chronic pain models^17^. Pathway activation scores were lower in PI16−/− mice as compared to WT mice (Figure S5). Moreover, the pathways neuropathic pain signaling in dorsal horn and PKA signaling pathway were inhibited in Pi16−/− mice, while they were activated in WT mice. The top 20 up-regulated genes after SNI included pain-related genes like GPR151, ATF3, GAL, CCKBR, NTS, and SEMA6A in both WT and Pi16−/− mice with reduced expression in Pi16−/− compared to WT. TGFβ was amongst the most prominent upstream regulators driving changes in gene expression. Overall pro-inflammatory cytokine gene expression was reduced in lumbar DRG of Pi16−/− animals post SNI as compared to WT-SNI (Figure S6).

Importantly, IPA disease and functions comparison analysis between WT and PI16−/− mice after SNI identified mainly biological functions (seven out of top ten) involved in immune cell migration and cell movement (Figure 7A). These included networks like leukocyte migration, inflammatory response, and quantity of cells and the corresponding genes associated with these networks included macrophage (ITGAM, CSF1R, Ccl9) and B/T cell (TNFRSF8/CD30, CD72) markers.

**Figure 7.**
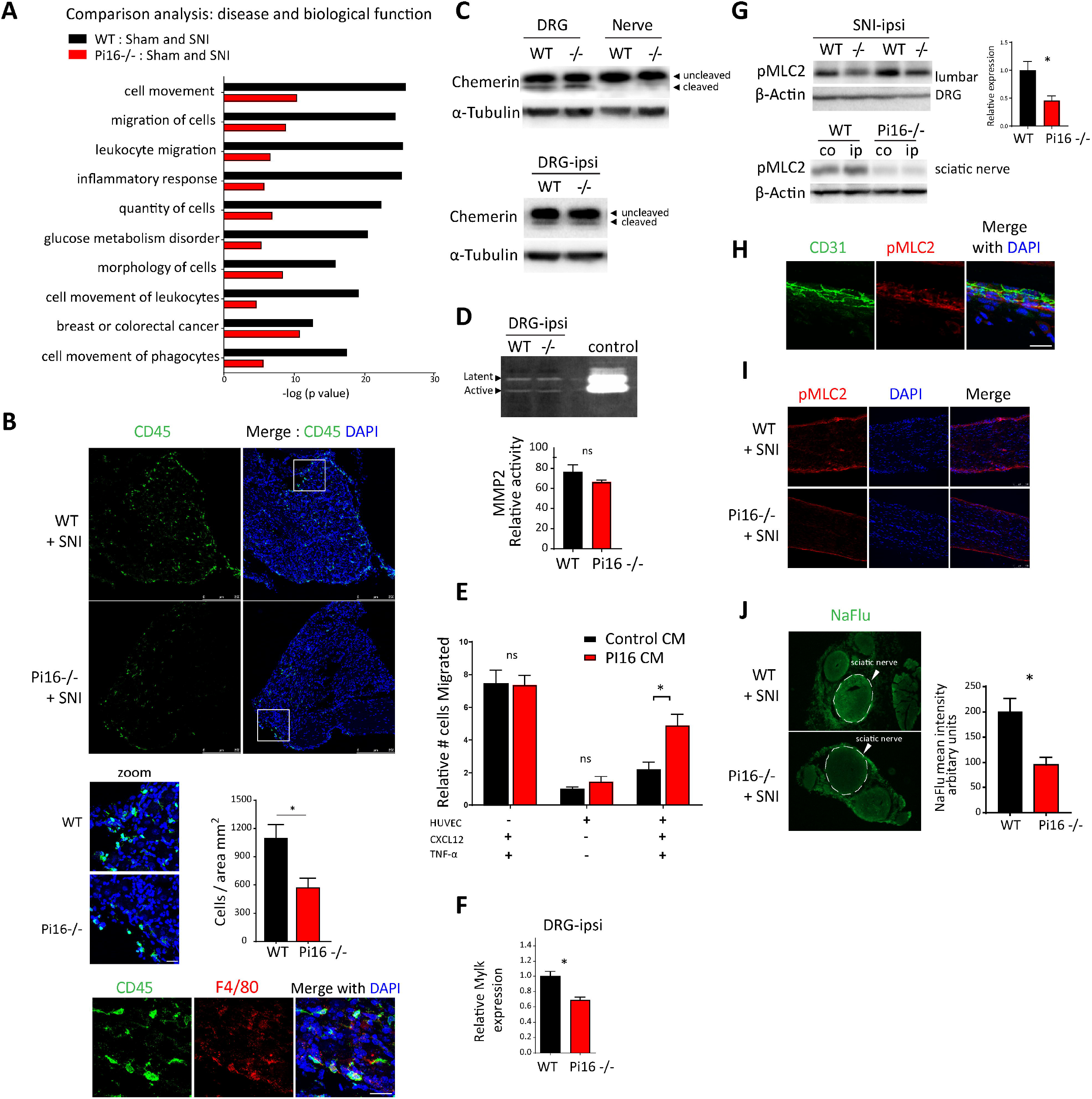
Pi16-deficiency inhibits the SNI-induced infiltration of leukocyte into the DRG and nerve. **(A)** IPA-based top ten disease biological functions different in lumbar DRG from WT and Pi16−/− mice post-SNI (day 9). IPA core analysis was performed between WT (WT-Sham and WT-SNI) and Pi16−/− groups (Pi16−/− Sham and Pi16−/− SNI) followed by comparison of the two core analyses. Black bars show change between WT-Sham and WT-SNI. Red bars show change between Pi16-Sham and Pi16-SNI. Y axis denotes –log (p value). (−0.5 < log2 Fold Change < 0.5), p 0.05. **(B)** Representative image of lumbar DRG ipsilateral to SNI showing immunostaining of leukocyte marker CD45 (green) and DAPI (blue) in WT and Pi16−/− mice on SNI day 5 (top panel). The zoom panel shows a magnified view of the white box (scale bar, 25 μm) and the bar graph shows quantitation data for CD45 staining from n=4 mice per group. T-test: *P < 0.05. Bottom panel shows staining of CD45 (green), macrophage marker F4/80 (red), and DAPI (blue), scale bar indicates 25 μm. **(C)** Western blot analysis of chemerin protein in naïve lumbar DRG and sciatic nerve of WT and Pi16−/− mice (upper panel) and in DRG from ipsilateral side after SNI in WT and Pi16−/− mice (lower panel). **(D)** Gelatin zymography for MMP-2 activity in the lumbar DRG of SNI WT and Pi16−/− mice. Bar graph shows quantitation data from n=3 mice per group. T-test: ns, not significant. **(E)** Effect of conditioned medium of PI16-overexpressing fibroblasts on monocyte migration across a TNF-α-activated endothelial cell monolayer in response to CXCL12 in the lower chamber either in control or PI16 conditioned medium (CM). Number of monocyte migrated to the lower chamber are plotted; relative to those of HUVEC (+) without activation (-TNF-α) and without chemoattractant (-CXCL12).*P < 0.05, two-way ANOVA, Turkey’s multiple comparisons test. ns, non-significant. **(F)** Real-time PCR analysis of Mylk expression in lumbar DRGs of WT and Pi16−/− mice. **P < 0.01 (analyzed using t test). **(G)** Upper panel-Western blot analysis of pMLC2 expression in ipsilateral lumbar DRGs after SNI. Blot showing pMLC2 levels in two separate WT and Pi16−/− mice and bar graph shows quantitation data. Bottom panel-Western blot analysis of pMLC2 level in sciatic nerve after SNI from contralateral (co) and ipsilateral (ip) side in WT and Pi16−/− mice. β-Actin was used as a loading control. (H) Representative images showing immunostaining of pMLC2 (red), DAPI (blue), and CD31 (green) in sciatic nerve longitudinal section in WT naïve mice. **(I)** Representative images showing immunostaining of pMLC2 (red) and DAPI (blue) in sciatic nerve longitudinal section from ipsilateral and contralateral side after SNI (day 5 post SNI). **(J)** Representative fluorescent image of sodium fluorescein (NaFlu) extravasation in sciatic nerve cross-section. WT and Pi16−/− mice received 40mg/kg of NaFlu i.v. and nerves were harvested 1hr later. Bar graph represents mean fluorescent intensity for NaFlu in the sciatic nerve (circle with dotted line) as determined from 3 mice per group. T-test: *P < 0.05.

To functionally validate these findings, we compared SNI-induced infiltration of leukocytes into the DRG and nerve between WT and Pi16−/− mice by immunostaining. Consistent with the transcriptome data, following SNI, Pi16−/− mice had lower numbers of CD45 positive cells in the lumbar DRG (Figure 7B) and nerve (Figure S7) than WT mice and the majority of these leukocytes stained positive for macrophage marker F4/80 (Figure 7B).

Previous studies have identified two possible targets of PI16 that could contribute to changes in leukocyte migration/infiltration, i.e. chemerin and MMP2^9,11^. PI16 has been described to inhibit the proteolytic activation of the chemoattractant chemerin^9,11^. However, we did not detect any difference in the expression of precursor- and activated-chemerin in the lumbar DRGs between WT and Pi16−/− post SNI (Figure 7C). The matrix metalloprotease MMP2 regulates leukocyte movement at the site of injury through regulation of chemokines^18,19^ and *in vitro* its activity can be regulated by PI16^11^. Gelatin zymography revealed similar MMP2 activity in WT and Pi16−/− lumbar DRGs post SNI mice (Figure 7D). These findings indicate that PI16 does not promote neuropathic pain via regulation of its putative targets MMP2 and chemerin.

To identify a potential direct effect of PI16 on movement of immune cells, we tested the transendothelial migration of monocytes in the presence of PI16 conditioned medium *in vitro.* Monocytes seeded in the top chamber were allowed to migrate across a monolayer of TNF-α-treated endothelial cells in response to chemotactic CXCL12 in the lower chamber. CXCL12 is a potent chemoattractant and our RNA sequencing data showed that it was increased in the DRG in response to SNI. In the presence of PI16 conditioned medium, monocyte migration across the endothelial barrier was increased (Figure 7E). We did not detect any effect of PI16-conditioned medium on migration in the absence of the endothelial barrier or in the absence of CXCL12.

Opening of the endothelial barrier is a crucial step allowing influx of leukocytes to the injured tissue. Our RNAseq data together with IPA analysis revealed lower levels of non-muscle Mylk, a well-established regulator of endothelial barrier, in DRGs from PI16−/− mice post-SNI compared to WT mice (Figure 7F). Mylk encodes myosin light chain kinase (MLCK) and MLCK-mediated phosphorylation of myosin light chain (MLC-2) promotes endothelial barrier permeability^20^. Notably, PI16−/− mice were protected against the SNI-induced increase in pMLC2 in ipsilateral DRGs and sciatic nerve (Figure 7G and I). MLC-2 is expressed by CD31-positive endothelial cells in close proximity to PI16 containing fibroblasts (Figure 7H).

To determine whether the reduced levels of pMLC2 in PI16−/− mice were associated with reduced endothelial permeability after SNI we injected the fluorescent tracer sodium fluorescein (NaFlu) intravenously. NaFlu accumulation was significantly lower in sciatic nerve of Pi16−/− nerve than in WT nerve (Figure 7J), indicating that Pi16 contributes to the SNI-induced increase in endothelial barrier permeability.

## Discussion

In this study, we identify for the first time the functional importance of the putative protease inhibitor PI16 in pain. We show that Pi16−/− mice are protected against mechanical allodynia in the SNI model of chronic neuropathic pain. In the DRG and peripheral nerve of naïve mice and mice after SNI, PI16 is not detectable in neurons, glia, leukocytes or endothelial cells, but was only detected in fibroblasts. Our data indicate that in vitro, PI16 enhances trans-endothelial migration of monocytes. In vivo, PI16−/− mice are protected against the SNI-induced increase in endothelial permeability and against the increase in leukocyte infiltration into the DRG and nerve. We propose that PI16 secretion by fibroblasts in DRG meninges and epi/perineurium represents a crucial permissive signal for establishment of chronic pain through promoting vascular permeability leukocyte infiltration. As a potential molecular mechanism we propose that PI16 promotes vascular permeability and cellular infiltration by increasing MLCK-dependent phosphorylation of the endothelial barrier regulator MLC2.

Until now, Pi16 had not been associated with DRGs, epi/perineurium, pain, or nerve injury and it’s in vivo function remained elusive. We identified Pi16 as a potential regulator of pain in an unbiased RNA screen comparing DRG samples from GRK2+/− mice who develop persistent pain and wild type mice who develop transient pain in response to local inflammation in the paw. We were interested in identifying genes which had not previously been associated with chronic pain. Mouse PI16 was first identified from a cardiac cDNA library and shown to be a glycosylated protein with expression in intercellular spaces, secretory vesicles, and a predicted role in the extracellular matrix^8^. Another study showed a strong upregulation of PI16 in a mouse model of heart failure and indicated a potential paracrine role of PI16 in cardiac hypertrophy^8,9^ However, genetic deletion of Pi16 conferred no phenotype at the level of cardiac structure or function indicating functional redundancy in the heart^9^. In our study, however, we demonstrate that PI16 is an important regulator of neuropathic pain. Mice with global Pi16 deletion are protected against neuropathic pain in SNI model (Figure 1). The protection against neuropathic pain in Pi16−/− mice was observed in both sexes, and is not a result of gross developmental structural abnormalities in DRG or nerve (Figure 1 and Figure S3). Thus, our findings strongly support a direct functional link between PI16 and neuropathic pain.

Our data indicate that along the pain neuroaxis, PI16 is mainly produced by fibroblasts in the meninges and in the perineurium. In accordance with our findings regarding PI16 expression profile in the DRG, published mouse single cell RNAseq data barely detected Pi16 mRNA in 23 out of 622 DRG neurons with reads below 100 in the majority of these 23 cells^21^. We show that SNI induces expansion of predominantly α-SMA-positive fibroblasts expressing PI16 in the meninges and epi/perineurium as well as at the injury site (Figure 2–4). Notably, SNI did not induce any detectable levels of PI16 in neuronal, glial, endothelial, and perineurial cells, or in endoneurial fibroblasts and infiltrating leukocytes in DRG and nerve. In addition, PI16 protein was not detectable in the spinal cord at baseline or after SNI. Moreover, although PI16 was detected in fibroblasts in the meninges of the spinal cord at baseline, we did not observe changes in distribution and expression upon SNI (Figure S4).

Fibroblasts play a key role in formation and maintenance of the extracellular matrix, but also actively secrete and respond to multiple cytokines, growth factors, and chemokines like IL-1, IL-6, TGF-β, TNF-α^22^. A previous study in rats suggested that fibroblasts in the meninges of the skull contribute to migraine pain through the production of pro-inflammatory cytokines like IL-6^23^. However, there was no direct evidence for a role of fibroblasts in this model of migraine pain. Our present findings imply a role of fibroblasts present in the meninges of the DRG and the perineurium of the sciatic nerve in controlling neuropathic pain through the production of PI16. Expansion of the perineurial layer has also been reported in the complete Freund’s adjuvant model of inflammatory pain in rats and we have preliminary evidence indicating that PI16 is increased in the epi/perineruium of mice with CFA-induced inflammatory pain. Expansion of the perineurium of peripheral nerves of humans with diabetic neuropathy has been reported, but the functional importance is unknown and PI16 has not been studied in this context^24,25^.

Our finding that PI16,aa protein that is not detectable in neurons and glia plays a key role in neuropathic pain add to the growing awareness that multiple non-neuronal cells types are actively involved in maintenance and resolution of chronic pain and may represent important targets for the treatment of chronic pain. In this respect we see PI16 as an interesting potentially druggable target for chronic pain considering its expression is restricted to very few organs (heart, bladder, mammary, fat) (https://tabula-muris.ds.czbiohub.org/)and, crucially, genetic deletion of Pi16 does not affect the cardiac function^9^, an organ in which PI16 is expressed at a high level.

PI16 is a secreted protein that may act as a paracrine factor on anatomically closely positioned endothelial cells in the DRG (Figure 2 and 6). We show that SNI induces expansion of fibroblasts expressing the myofibroblast marker α-SMA in the meninges that is associated with an increase in PI16 expression as seen by Western blots and immunostaining (Figure 3 and 4). In addition, in vitro differentiation of fibroblasts into myofibroblasts in response to TGFβ1, led to an upregulation and secretion of PI16 from primary fibroblasts (Figure 6). Komuta et al. 2010 showed TGF-β1 receptor increases in the meninges following brain injury and TGF-β1 is involved in activating meningeal fibroblasts^26^. Moreover, we found TGF-β1 mRNA was the primary transcriptional regulator driving changes in gene expression in response to SNI in our RNA-seq data and this could be how PI16 is upregulated as a result of SNI. Notably, PI16 positive fibroblasts remain restricted at the DRG borders in the meninges after SNI and did not invade the neuronal parenchyma of the DRG, making it unlikely that PI16 directly targets nociceptors.

A combination of transcriptomic, immunofluorescence, and cell culture studies employed here revealed that PI16 promotes migration of immune cells across the endothelial barrier. In vivo, deficiency of PI16 caused reduced infiltration of leukocytes in DRG as tested on day 5 after SNI (Figure 7) and at this time point Pi16−/− mice were protected against SNI-induced pain (Figure 1). Conversely, in vitro PI16 conditioned medium promotes migration of immune cells in a transendothelial migration assay (Figure 7). RNA-seq with lumbar DRGs from SNI-treated mice revealed multiple cellular infiltration and migration pathways downregulated in Pi16−/− mice (Figure 7). Genes involved in these functional networks were mostly cytokines (e.g. CRH, CSF1, CXCL14, Ccl9, TIMP1, IL1RN, CCL2, SPP1, IL6); growth factors (e.g. FGF3, GRP, IGF1, ANGPT1, NOG); GPCRs (MC4R, GPR34, CYSLTR1, CCR5, CCR2, CCR1, CX3CR1, C3AR1); extracellular proteins (e.g. SERPINE1, LGALS3, TNC, COCH, LGALS1, OLFM4, IGFBP3, CFH); signaling kinases (e.g. CAMK1, PIK3CG, MYLK, CDKN1A, EGFR, CSF1R, DDR2, TGFBR1) and transmembrane receptors (e.g. TNFRSF8, MSR1, FCER1G, TLR7, PLAUR, IL17RA, IL1R1, CD72, LILRB3, CR2, TREM2, ITGAM, CXADR, UNC5B). A representation of genes from different protein families reflects a multilevel impact of reduced cellular infiltration caused by PI16 deficiency. It remains to be determined which of these occur downstream of changes in leukocyte infiltration in PI16−/− mice.

How does PI16 influence the migration of leukocytes into the DRG? PI16-dependent regulation of chemerin and MMP2 has been proposed and these proteins can regulate the movement of immune cells and overall inflammatory load in tissues^9,11^. However, we did not detect changes in the level of processed chemrin or MMP2 when comparing samples from WT and PI16−/− mice under naïve conditions or after SNI (Figure 7). An increase in the permeability of endothelial barrier is essential for the influx of leukocytes in to the tissues following nerve injury. Lim et al., (2014) demonstrated the breakdown of the blood nerve barrier following nerve injury drives neuropathic pain by facilitating extravasation of immune cells into the nerve^27^. Our RNAseq data following SNI indicated an interaction between PI16 and MLCK, a kinase involved in breakdown of endothelial barrier. We found in response to SNI Pi16−/− mice have reduced Mlck expression and correspondingly, repressed phospho-MLC2 activation in DRG and sciatic nerve. MLCK and its substrate MLC2 maintain integrity of endothelial barrier and their function has been highlighted in diseased models of lung injury, pancreatitis, and atherosclerosis^20,28–30^. In a mouse model of lung injury in response to endotoxin lipopolysaccharide, MLCK−/− mice preserve endothelial barrier function compared to wildtype and show reduced leukocyte infiltration in lungs^31^. Consistent with reduced pMLC-2 levels, Pi16−/− mice show reduced cellular infiltration and vascular permeability as measured by sodium fluorescein dye. The specific mechanism how PI16 modulates MLCK/MLC-2 pathway in endothelial cells remains to be established.

Taken together, our findings support a model in which nerve injury enhances production and secretion of PI16 by meningeal and epi/perineurial fibroblasts which promotes neuropathic pain by increasing the permeability of the endothelial barrier and facilitating immune cell infiltration into DRG and nerve. We also propose that PI16 mediated control of MLCK-dependent phosphorylation of MLC-2 contributes to the increased permeability of the endothelial barrier in neuropathic pain. In view of the limited cellular and organ distribution of PI16 we propose that PI16 represents an attractive novel potential target for pain management that is unlikely to have abuse liability.

## Acknowledgments

This work was supported by the National Institute of Neurological Disorders and Stroke of the National Institutes of Health RO1 NS073939 and RO1 NS074999. We acknowledge Kenneth Dunner Jr. at the High Resolution Electron Microscopy Facility at MD Anderson Cancer Center for tissue processing and image capturing. We thank Ms. Itee Mahant, Dr. Angie C.A. Chiang, Ms. Jenolyn Alexander, and Mr. Jules D Edralin for technical help.

## Author Contributions

AK and CJH designed experiments, supervised data analysis, and edited the manuscript. PS designed and performed experiments, analyzed the data, and wrote the manuscript. JM, XJH, RT performed experiments. All authors approved the final manuscript. BP contributed to RNA sequencing analysis.

## Footnotes

We have no conflicts of interests related to this study.

## Corresponding author

Correspondence to Prof. Annemieke Kavelaars

## Materials and Methods

### Animals

Male and female B6:129S mice homozygous for global Pi16 deletion (MMRRC stock number 032520-UCD) and their wild type control littermates at an age of 8-12 weeks were used in the study. All procedures were performed following the ARRIVE guidelines, and are in accordance with National Institutes of Health Guidelines for the Care and Use of Laboratory Animals and the Ethical Issues of the International Association for the Study of Pain^32^. Studies were approved by the Institutional Animal Care and Use Committee of the University of Texas MD Anderson Cancer Center. Human DRGs used for RT-PCR were collected from patient donors at MD Anderson Cancer Center who had provided legal written consent. The protocol was reviewed and approved by the MD Anderson Cancer Center Institutional Review Board and the donors were undergoing spinal surgery for disease treatment wherein a spinal nerve root was sacrificed as the standard of care. All pain measures and immunofluorescence analyses were performed by investigators blinded to treatment.

### Spared nerve injury model

SNI surgery was performed as described previously^33,34^. Briefly, mice were anesthetized under isoflurane and the three peripheral branches of the left sciatic nerve (sural, common peroneal and tibial) were exposed. The common peroneal and tibial nerves are tightly ligated and then transected distal to the ligation. The sural nerve was kept intact. For sham surgery, nerves were exposed but not ligated or transected. Von Frey measurements were performed starting from 4 days after SNI surgery. All biochemical and immunofluorescence assays were done at an early (d4-d8) and/or late (d22-d45) time point post SNI.

### Measurement of mechanical allodynia

Mechanical allodynia was measured as the hind paw withdrawal response to von Frey hair stimulation and the 50% paw withdrawal threshold was calculated using the up-and-down method^35^. Decrease in paw-withdrawal threshold is indicative of mechanical allodynia. In brief, animals were placed on a wire-grid base, through which the von Frey hairs (Stoelting, Wood Dale, IL, USA) with bending force range from 0.02 to 1.4 g were applied. Clear paw withdrawal, shaking, or licking was considered as nociceptive-like responses.

### Cell culture and reagents

Mouse perineurial fibroblasts (M1710-57, ScienCell Research Laboratories) were cultured in complete fibroblast medium (ScienCell Research Laboratories catalog number 2301) per the manufacturer’s instructions. For differentiation of fibroblasts to myofibroblasts 5ng/ml of TGF-β1 (7666-MB, R&D Systems) was used. HUVEC cell (PromoCell C-12203) were cultured in Endothelial Cell Growth medium (PromoCell C-22210) supplemented with Supplmix (C-39215). L cells (ATCC^®^ CRL-2648) were cultured in high glucose Dulbecco’s modified Eagle medium (GE Healthcare, Piscataway, NJ, USA) plus 10% fetal bovine serum (Gibco, Carlsbad, CA, USA). L cells were transiently transfected with Pi16-tGFP (Origne MG219996) and pm-Turq-ER (Addgene 36204) with Lipofectamine 2000 (Invitrogen, Grand Island, NY, USA) according to the manufacturer’s instructions. To prepare conditioned medium, L cells were seeded in 6-well plates and incubated overnight, followed by transfection with 1.5 μg of Pi16-tGFP plasmid. The medium was replaced with serum free DMEM and transfected cells were incubated for 48 hours. Cell supernatant was collected from culture and centrifuged to pellet debris and dead cells and filtered through a 0.2μm filter. Supernatant from untransfected or Pi16-tGFP transfected cells was diluted with fresh medium (1 part of fresh medium to 2 parts of supernatant) to get control conditioned medium and Pi16 conditioned medium) respectively.

### Real-time quantitative PCR

The first-strand cDNA was synthesized using one-to-two μg of RNA using high capacity cDNA reverse transcription kit (Applied Biosystems) and qPCR was performed on an Applied Biosystems ViiA using Taqman primers (Integrated DNA Technologies, Coraville, IA, USA). Relative quantitative measurement of target gene levels corrected for GAPDH or β-actin was performed using the ΔΔCt method. Primer used were as follows: Human PI16 (IDT, Hs.PT.58.39664750), Mouse Pi16 (Mm.PT.58.8467939 and Mm.PT.58.22010817).

### RNA extraction, library preparation for sequencing, and RNA-seq Data analyses

Transcriptional changes in the lumbar DRGs were investigated using whole-genome RNA sequencing by the RNA Sequencing Core Lab at MD Anderson Cancer Center. RNA-seq was performed on triplicate samples and the RIN number for all samples was >7. Mice were perfused intracardially with chilled PBS and lumbar DRGs (L3–L5) were collected without desheathing the meninges. DRGs were collected 5 hours after the PGE2 or vehicle treatment from WT and GRK2+/− mice and 9 days post SNI or Sham surgery from WT and Pi16−/− mice. DRGs were homogenized in TRIzol reagent (Invitrogen, Carlsbad, CA, USA) and total RNA was extracted with an RNeasy MinElute Cleanup Kit (Qiagen, Hilden, Germany). For RNA-seq from GRK2+/− mice, DRGs were pooled from two mice to get one sample.

The sample libraries were generated using the Stranded mRNA-Seq kit (Kapa Biosystems, Wilmington, MA) following the manufacturer’s guidelines. A 75-nt paired-end run format was performed using a HiSeq 4000 Sequencer as previously described^36^. Six samples were sequenced in a single lane. RNA sequencing data analysis was performed using a comprehensive in-house RNASeq pipeline, which used STAR to align paired-end reads to the mouse reference genome (mm10 version) and featureCounts to obtain expression counts of genes. The quality of raw and aligned reads was assessed using FastQC, RSeQC, and qualimap. In both GRK2+/− and Pi16−/− analyses, we used principal component analysis and unsupervised hierarchical clustering to analyze and samples and R Bioconductor packages EdgeR and DESeq2 to compare expressions of genes between groups of interest. We excluded genes with on average less than 10 reads per samples and selected genes with adjusted p-value < 0.05 as differentially expressed. For SNI dataset, further analysis was performed with Ingenuity Pathway Analysis (IPA; Qiagen Inc., https://www.qiagenbioinformatics.com/products/ingenuity-pathway-analysis/) to identify biological processes and canonical pathways that were potentially affected by PI16 deficiency. A data set of differentially expressed genes (adjusted p<0.05) containing gene identifiers, their corresponding expression values (log_2_Ratio), and P-values was uploaded on IPA for core analysis.

### Immunofluorescence and microscopy

DRG (lumbar L3-L5), spinal cord, and sciatic nerve were collected from mice perfused with PBS followed by 4% paraformaldehyde in PBS. Spinal cord meninges were gently peeled with forceps from the vertebrae in a single layer. Tissues were cryo-protected in 20% sucrose and embedded and frozen in 1:1 mix of optimal cutting temperature compound (OCT; Bayer Corporation) and 20% sucrose, and sliced into 14 μm sections. Sections were permeabilized with 0.3% Triton-x-100 and blocked with normal serum (5%), saponin (0.1%), and BSA (2%) and stained with following primary antibodies: PI16 (1:75, R&D Systems AF4929), GLUT1 (1:250, Abcam ab652), α-SMA (1:100, Abcam ab5694), P4HB (1:250, Abcam ab137110), IBA1 (1:300, Wako 019-19741), NFH (1:200, Millipore AB1989), CGRP (1:200, Abcam ab36001), IB4 (1:100, Vector Labs B-1205), CD31 (1:50, Abcam Ab28364), CLDN1 (1:250, Abcam ab15098), Collagen IV (1:100, Southern Biotech 1340-01), PGP9.5 (1:500, Abcam ab108986), P4HB (Abcam ab137110), CD45 (1:50, BD Biosciences 550539), pMLC2 (1:200, Cell Signaling 3675S) followed by Alexa-594, Alexa-488, or Alexa-647 conjugated secondary donkey or goat antibodies (Invitrogen, Grand Island, NY). Nuclei was stained with DAPI and for negative control, primary antibody was omitted. Sections were visualized using a Leica SPE DMI 4000B confocal microscope or EVOS fluorescence microscope. The mean intensity of fluorescence and the percent area positive were calculated using Leica or NIH ImageJ software. For TEM analysis of mitochondrial morphology, nerves were fixed in 2% glutaraldehyde plus 2% PFA in PBS and processed as previously described^37^.

### Western blotting

Tissue and cell lysates were prepared using Bioruptor (Diagenode) in RIPA buffer (10 mM Tris-HCl, pH 6.8, 100 mM NaCl, 1 mM EDTA, 10% glycerol, 1% Triton X-100, 0.1% SDS, 0.5% sodium deoxycholate, 2 mM Na3VO4, 50 mM NaF) supplemented with protease inhibitor cocktail and phosphatase inhibitors. Protein content was assessed using Bradford (Bio-Rad Laboratories, Hercules, CA, USA) or BCA assay (Pierce, Rockford, IL, USA). Equivalent amounts of protein or equal volume from each sample were run on SDS-polyacrylamide gels and transferred to polyvinylidene difluoride membranes (GE Healthcare, Piscataway, NJ, USA). Following blocking in 5% milk or bovine serum albumin the blots were incubated with primary antibodies and analyzed by HRP-conjugated secondary antibodies (Jackson Laboratories). The following primary antibodies were used in the study: PI16 (1:750, R&D Systems AF4929), α-SMA (1:1500, Abcam ab5694), β-Actin (1:50,000, Sigma A3854), α-Tubulin (1:2000, Cell Signaling 2144S), pMLC2 (Cell Signaling 3674S, 3675S), Chemerin (R&D Systems AF2325). For detection of signal, enhanced chemiluminescent agent (GE Healthcare, Piscataway, NJ, USA) or Clarity Western ECL Substrate (Bio-Rad, Hercules, CA, USA) was used followed by imaging on LAS4000 chemiluminescence system. Band density was determined using LAS Image Quant software (GE Healthcare, Piscataway, NJ, USA).

### Sodium fluorescein permeability assay

The permeability of vascular endothelium in nerve was assayed by extravasation of the fluorescent dye sodium fluorescein (Sigma Aldrich, St. Louis, MO)^27^. Sodium fluorescein (40mg/kg) was injected through the tail vein on day5 post SNI surgery and allowed to circulate for 1 hour. Mice were perfused with ice-cold PBS with heparin intracardially and nerves were collected and processed as described above in immunofluorescence section. Cross-sections of the nerve were quantified by EVOS fluorescence microscope and the mean intensity of fluorescence and the percent area positive were calculated using NIH ImageJ software.

### Transwell assay

Transwell inserts (Corning 6.5mm diameter insert, 5um pore size, PET membrane, 24 well plate) were coated with collagen I (30ug/ml overnight, Gibco A10483-01) and fibronectin (7.5ug/ml for 1 hour, Corning 354008). HUVEC cells (2.0 × 10^4^, PromoCell C-12203) were seeded in endothelial medium (PromoCell C-22210) in the inserts and grown till a confluent monolayer of cells was formed (3 days). In parallel, L cells were transfected with PI16-tGFP construct and PI16 conditioned medium was prepared. In addition, THP-1 monocytes were cultured in RPMI medium and treated with phorbol 12-myristate 13-acetate (PMA) (300 nM, 48 h) to differentiate them into monocytes. On the day of assay, HUVEC cells were activated with 500 units/ml of TNFα (210-TA-005) either in control or PI16 conditioned medium following by addition of 1 × 10^5^ THP-1 monocytes to the top chamber. Human CXCL12 (SDF-1α, 300-28A, 50ng/ml) either in control or PI16 conditioned medium was added to the bottom chamber. Cells were allowed to migrate for 5 hours. Inserts were washed with PBS followed by fixation with 4% PFA and staining with 0.2% crystal violet. Cells on the top of the insert were removed using a cotton swab and only cells which migrated across the membrane were counted under a microscope.

### Statistics

Data are expressed as mean ± S.E.M. Data were analyzed using GraphPad Prism 6. Statistical analyses were carried out using Student’s t test, one-way or two-way ANOVA with or without repeated measure followed by Turkey or Bonferroni analysis. A p value less than 0.05 was considered significant.

