## Supplementary Figures S1-S7 for "PI16 is a non-neuronal regulator of neuropathic pain"

**A**

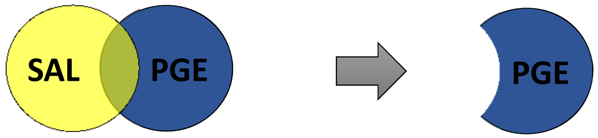


| ID | gene | WT-SAL | WT-PGE | KO-SAL | KO-PGE | PGE (WT->KO) regulated | PGE (WT->KO) q_value | SAL (WT->KO) regulated | SAL (WT->KO) q_value | WT (SAL->PGE) regulated | WT (SAL->PGE) q_value | KO (SAL->PGE) regulated | KO (SAL->PGE) q_value |
|---|---|---|---|---|---|---|---|---|---|---|---|---|---|
| **10906** | Iigp1 | 8.66675 | 11.2197 | 9.26582 | 5.81304 | down | 0.012525 | no | 0.998727 | up | 0.028157 | down | 0.000762 |
| **2948** | Gm12250 | 1.38477 | 2.18125 | 1.62094 | 1.20611 | down | 0.012525 | no | 0.998727 | up | 0.011328 | no | 0.180398 |
| **14859** | Gbp2 | 16.3362 | 19.8469 | 16.7203 | 13.3295 | down | 0.012525 | no | 0.998727 | no | 0.090334 | down | 0.041924 |
| **2950** | Irgm2 | 8.26463 | 10.2182 | 7.94863 | 6.98765 | down | 0.012525 | no | 0.998727 | no | 0.065261 | no | 0.368016 |
| **9677** | Pi16 | 22.6126 | 15.9402 | 23.4075 | 21.9794 | up | 0.012525 | no | 0.998727 | down | 0.001487 | no | 0.714363 |

**B**

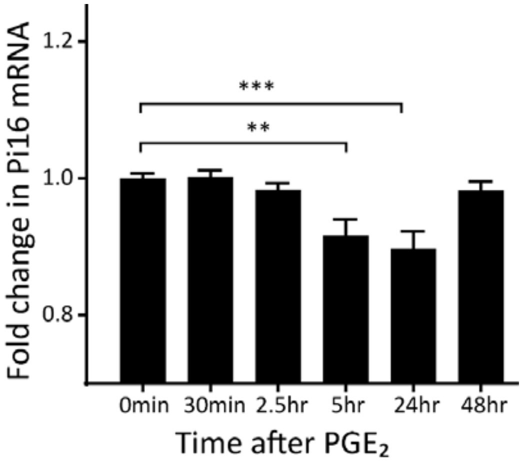


**Figure S1, Related to Figure 1. (A)** Normalized log-counts-per-million reads for novel differentially expressed genes (DEG) between WT and GRK2+/- (KO) mice associated with pain sensitivity. DEG were identified for the SAL group (between WT-SAL and KO-SAL) and $PGE_2$ group (between WT-$PGE_2$ and KO-$PGE_2$) and the overlapping genes between the two groups were eliminated (i.e. genes that differed due to genotype). **(B)** Validation of PGE2-induced changes in PI16 mRNA by real-time PCR analysis of PI16 expression in independent samples of lumbar DRGs of WT mice at various time points after $PGE_2$ treatment (100 ng/paw). GAPDH was used for normalization. n = 3 mice per time point; One-way ANOVA followed by post hoc (Turkey) test: **P < 0.005, ***P < 0.0005.

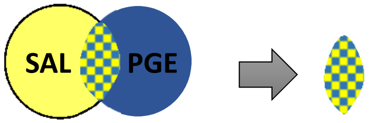


| ID | gene | WT-PGE | KO-PGE | WT-SAL | KO-SAL | SAL (WT->KO) regulated | SAL (WT->KO) q_value |
|---|---|---|---|---|---|---|---|
| 450 | Tnni1 | 12.8927 | 4.68409 | 25.3463 | 5.53242 | down | 0.007541 |
| 6744 | Tnnc1 | 23.2529 | 6.82758 | 52.729 | 9.62632 | down | 0.007541 |
| 7399 | Myh7 | 3.59527 | 0.937171 | 6.78397 | 0.836385 | down | 0.007541 |
| 8325 | Mb | 60.1451 | 41.3157 | 116.967 | 50.5631 | down | 0.007541 |
| 11727 | Adrbk1 (GRK2) | 69.8604 | 42.893 | 66.8315 | 38.5189 | down | 0.007541 |
| 15435 | Myoz2 | 5.92071 | 2.4654 | 9.87762 | 4.5512 | down | 0.007541 |
| 17704 | Myl2 | 41.355 | 9.54716 | 79.603 | 16.624 | down | 0.007541 |
| 21238 | Tnnt1 | 16.0347 | 4.00184 | 31.1835 | 6.06305 | down | 0.007541 |
| 24130 | Sln | 7.71582 | 1.69838 | 18.1172 | 2.15711 | down | 0.007541 |
| 24501 | Myl3 | 17.2083 | 4.33783 | 28.6223 | 5.24341 | down | 0.007541 |
| 1782 | Prtn3 | 18.1083 | 27.3485 | 26.9032 | 19.0768 | down | 0.013198 |
| 1783 | Elane | 18.6586 | 27.7646 | 27.8844 | 18.3039 | down | 0.007541 |
| 1785 | Cfd | 80.721 | 125.635 | 145.331 | 94.0642 | down | 0.007541 |
| 3035 | Myh1 | 2.30788 | 5.42612 | 5.90265 | 3.5096 | down | 0.007541 |
| 3106 | Eno3 | 25.929 | 41.5637 | 71.2126 | 49.1519 | down | 0.007541 |
| 3251 | Slfn4 | 2.22643 | 3.47022 | 3.78227 | 2.30737 | down | 0.007541 |
| 3303 | Mpo | 10.8255 | 16.3733 | 15.8396 | 10.5542 | down | 0.007541 |
| 3681 | Pgam2 | 11.9112 | 18.978 | 32.4448 | 18.0839 | down | 0.007541 |
| 4418 | Slc4a1 | 22.3193 | 29.3149 | 27.9448 | 20.391 | down | 0.022787 |
| 6491 | Cmya5 | 1.80324 | 2.47959 | 4.10177 | 2.98727 | down | 0.018415 |
| 7193 | Myoz1 | 6.85066 | 13.6969 | 27.3086 | 18.8787 | down | 0.031464 |
| 8750 | Adipoq | 13.1821 | 19.9494 | 22.9245 | 15.4039 | down | 0.007541 |
| 11739 | Actn3 | 10.1665 | 18.9589 | 40.4975 | 27.5767 | down | 0.026934 |
| 13413 | Lcn2 | 107.925 | 154.006 | 157.345 | 116.103 | down | 0.013198 |
| 13568 | Ttn | 0.451306 | 0.621851 | 0.900694 | 0.510824 | down | 0.007541 |
| 16436 | Tpm2 | 23.2742 | 32.4952 | 63.7371 | 35.3539 | down | 0.007541 |
| 17551 | Ibsp | 9.26908 | 13.3032 | 17.4578 | 12.1261 | down | 0.031464 |
| 18268 | Igj | 7.20812 | 11.2176 | 9.26268 | 4.50295 | down | 0.007541 |
| 19860 | Cidec | 8.9779 | 13.318 | 13.6596 | 8.06286 | down | 0.007541 |
| 20270 | Ckm | 66.6505 | 121.217 | 229.849 | 157.882 | down | 0.007541 |
| 21502 | Cd177 | 6.71992 | 9.94331 | 8.70326 | 5.44563 | down | 0.007541 |
| 23661 | Hp | 26.482 | 40.0049 | 38.1018 | 25.0785 | down | 0.007541 |
| 23782 | Acta1 | 104.288 | 145.984 | 277.514 | 167.528 | down | 0.007541 |
| 24497 | Ngp | 230.858 | 323.462 | 307.812 | 228.971 | down | 0.040608 |
| 24511 | Ltf | 38.6149 | 55.1005 | 49.2932 | 32.4304 | down | 0.007541 |
| 25885 | Alas2 | 52.6075 | 70.2154 | 80.5557 | 59.0525 | down | 0.007541 |
| 7617 | Dct | 0.359117 | 1.34026 | 0.744143 | 1.92369 | up | 0.007541 |
| 15784 | Tyrp1 | 5.12758 | 11.0774 | 6.51297 | 11.872 | up | 0.007541 |
| 24855 | Zbtb16 | 1.284 | 2.39998 | 1.03602 | 2.48913 | up | 0.007541 |

**Figure S2, Related to Figure 1.** Normalized log-counts-per-million reads for DEG that differed due to GRK2+/- genotype between WT-SAL and KO-SAL.

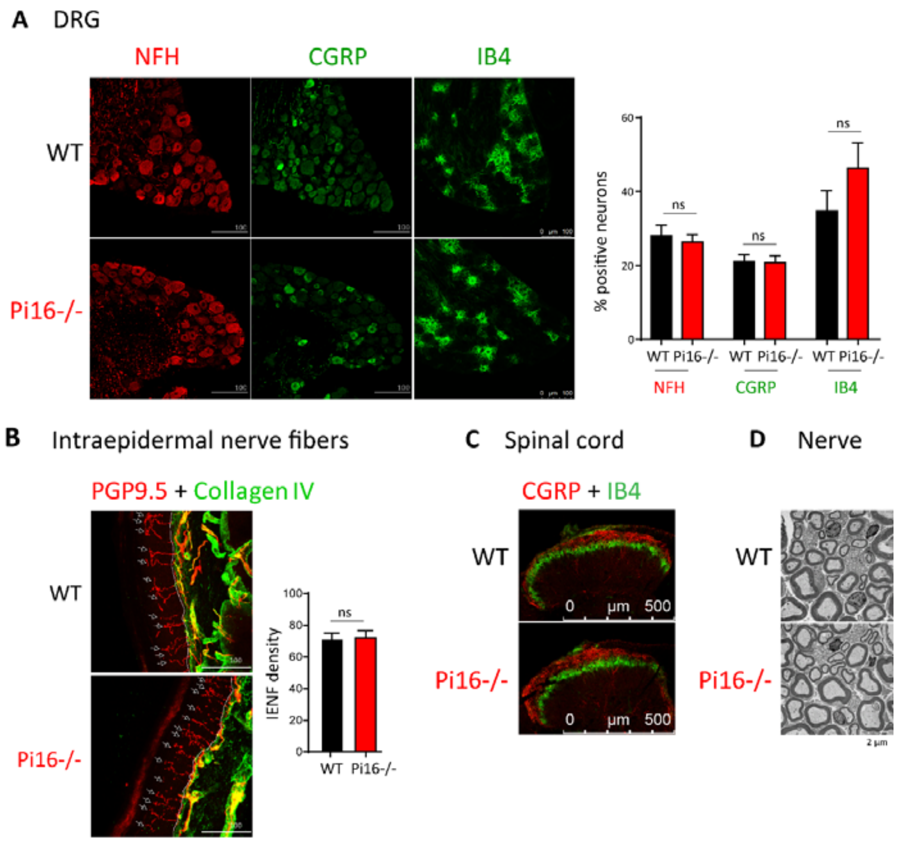


**Figure S3, Related to Figure 1.** Pi16-/- knockout mice have normal distribution pattern of neuronal subpoupulations. **(A)** Representative images showing lumbar DRG rom WT and Pi16-/- mice stained for NF200 A-fiber neurons (NFH), CGRP peptidergic neurons, and IB4 nonpeptidergic neurons in lumbar-DRGs Scale bar indicates 100 µm. Bar graph represents percent of neurons staining positive for each marker. No significant differences were observed between WT and Pi16-/- mice. **(B)** Immunostaining with pan-neuronal marker PGP9.5 (red) and basement collagen membrane (green) to identify IENFs in hind paw skin sections from WT and Pi16-/- mice. Representative confocal images show IENF (white arrows) crossing the basement collagen membrane. Bar graph shows IENF density expressed as number of nerve fibers crossing the basement membrane/length of the basement membrane. Scale bar indicates 100 µm. **(C)** Representative image of lumbar spinal cord of WT and Pi16-/- mice stained for CGRP peptidergic neurons and IB4 nonpeptidergic neurons. **(D)** Electron micrograph of a sectioned sciatic nerve from WT and Pi16-/- mice showing normal myelin wrapping and nerve fiber structure. Scale bar indicates 2 µm.

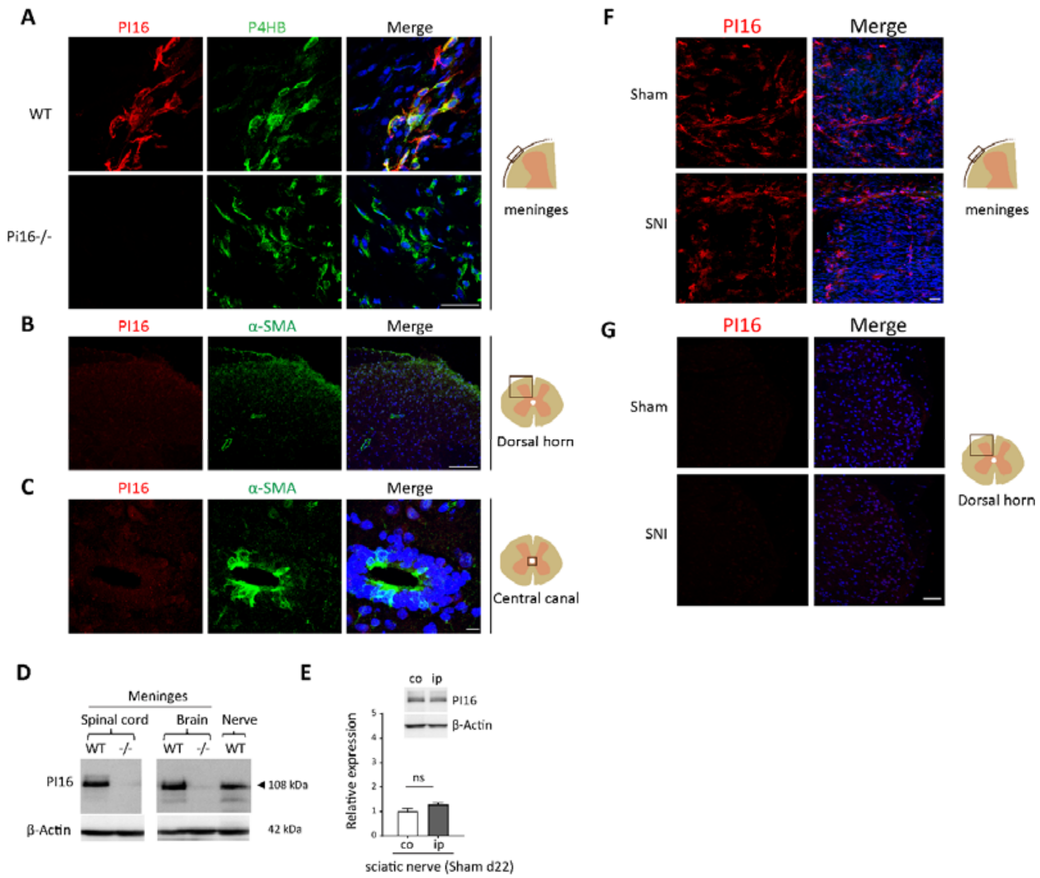


**Figure S4, Related to Figure 2-4.** PI16 is expressed in fibroblasts in the meninges of spinal cord and SNI does not induce changes in spinal cord PI16 expression. **(A)** Representative image of spinal cord meninges (from naïve WT and Pi16-/- mice stained with PI16 (red) and fibroblast marker P4HB (green). **(B) and (C)** Representative spinal cord section from naïve WT mice showing dorsal horn area, B and central canal, C stained with PI16 (red), α-SMA (green), and DAPI (blue). Scale bar indicates 50 µm for A, 100 µm for B, and 10 µm for C. **(D)** Western blot analysis showing PI16 protein expression (detected as a 108kDa band) in spinal cord and brain meninges from naïve animals. **(E)** Western blot analysis of PI16 protein expression in ipsilateral (ip) and contralateral (co) sciatic nerve at 22 days after Sham surgery (t test, ns not significant). **(F)** Representative image showing PI16 (red) and DAPI (blue) staining in spinal cord meninges in WT mice after Sham or SNI surgery (day 5). **(G)** Representative image of spinal cord section showing dorsal horn of ipsilateral side stained with PI16 (red) and DAPI (blue). Scale bar for B and C indicates 50 µm.

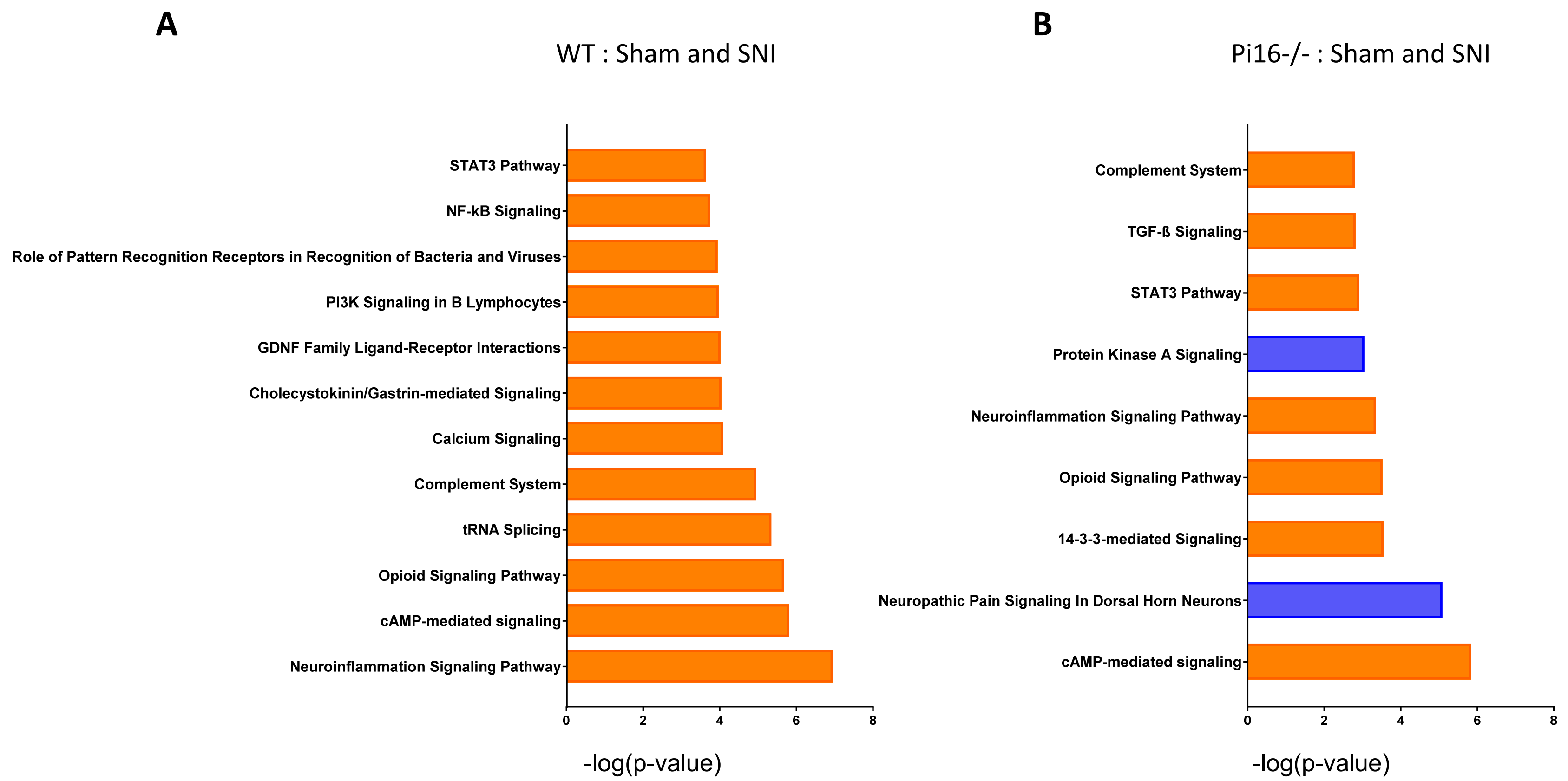


**Figure S5, Related to Figure 7.** Top IPA canonical pathways assigned to differentially regulated genes between **(A)** WT + Sham and WT + SNI for 956 DEG with a cutoff (−0.4 < log2 Fold Change < 0.4), p 0.05 and **(B)** Pi16 + Sham and Pi16 + SNI for 1138 DEG with a cutoff (−0.2 < log2 Fold Change < 0.2) and p  0.05. $\log_2$ Fold Change cutoff was adjusted to normalize the number of genes used for annotating canonical pathways.

**WT : Sham vs SNI**

| Upstream Regulator | Expr Log Ratio | Activation z-score | p-value of overlap | Target molecules in dataset |
|---|---|---|---|---|
| DKK3 | -0.325 | | 0.0142 | ANGPT1,TAGLN |
| TNFSF11 | 0.16 | 3.937 | 5.83E-12 | ADGRE1,AIF1,AQP9,ATP6V0D2,C3AR1,CCL2,Ccl8,Ccl9,CCR1,CDKN1A,CSF1R,CTSK,IFRD1,IGF1,IL13RA1,IL1RN,IL6,ITGAM,JUN,NFIL3,NFKBIZ, PLAUR,PRDM1, SDC1,SERPINE1,SPP1,SRXN1 |
| IL24 | 0.237 | 1.204 | 0.0143 | CASP3,GADD45A,GADD45G,PRDM1 |
| IL7 | 0.341 | 2.42 | 0.000823 | BTLA,CASP3,CCL2,CDKN1A,CSF1,CTSK,HAVCR2,IL1R1,IL6,Ly6a (includes others) |
| IL33 | 0.372 | 1.729 | 0.00337 | ABCA1,CCR2,IL6,ITGAM,LEPR,MSR1,TIMP1 |
| EDN1 | 0.376 | 2.985 | 0.0000182 | ANXA1,CCL2,CYBB,EGFR,FST,IL6,ITGAM,JUN,LEPR,ODC1,PLAUR,SDC1,SERPINE1, TIMP1,TNC |
| CYTL1 | 0.399 | | 0.0209 | IGF1 |
| CXCL12 | 0.424 | 2.354 | 0.000822 | AIF1,CCL2,CTSK,EGFR,IL6,JUN,LOXL2,Ly6a (includes others),PPEF1,RHOC,TIMP1,TNNC1 |
| IL1B | 0.451 | 4.774 | 8.15E-17 | ADAM8,AIF1,ANGPT1,ANXA1,ATF3,BAMBI,CASP3,CCKAR,CCL2,Ccl9,CCR1,CCR2,CCR5,CDKN1A,CREM,CRH,CSF1,CTSS,CX3CR1,CYBB, CYSLTR1,F13A1,FGF7,FST,GADD45A,HPGDS,Hrk,IBSP,IFI16,IFRD1,IGF1,IGFBP3,IL1R1,IL1RN,IL6,ITGAM,JUN,LAMB3,LOX,MAFA,MYLK, NFIL3,NFKBIZ,NPY,NTRK1,ODC1,PAPPA,PDE4B,PHLDA1,PTHLH,S100A10,SDC1,SERPINE1,SLC6A4,SPP1,STAR,STMN2,TIMP1,TLR7,TLR8,TREM2 |
| Ccl6 | 0.466 | | 0.0142 | CTSK,CTSS |
| IL6 | 0.503 | 4.549 | 6.47E-18 | ABCA1,ACVR1,ADGRE1,ANXA1,ATF3,CASP3,CCL2,CCR1,CCR2,CCR5,CD163,CD38,CD53,CD68,CDKN1A,CDKN3,CRH,CRLF1,CSF1,CTSC,CTSK, CX3CR1,CYBB,CYP1B1, EGFR,FCGR1A,GADD45A,GADD45G,GAL,GAP43,HPGD,IFI16,Ifi202b,IGF1, IGFBP3,IL1R1,IL1RN,IL6,ITGAM, ITPR1,JUN,KIF22,LGALS1,MSR1,NPY,PTPRC, REG1A,SERPINE1,SPP1,STEAP4,TH,TIMP1,TLR1,TLR7,TLR8,TNC,TNFRSF12A |
| SPP1 | 0.515 | 2.634 | 0.0000255 | ANGPT1,CCL2,CD163,CDKN1A,DSP,EGFR,IGF1,IL6,LOX,SERPINE1,SNAI2,SPP1, TGFBR1,TIMP1 |
| CCL2 | 0.552 | 1.233 | 0.00017 | ABCA1,CCL2,CCR2,HDC,IGF1,IL6,SERPINE1,TIMP1 |
| IL1RN | 0.597 | -1.637 | 0.000000448 | ATF3,CCL2,CHAC1,CRH,CTSS,IGF1,IL6,KLF6,NTRK1,SERPINE1,SLC15A3, SLC6A4,SPP1,TIMP1 |
| TIMP1 | 0.607 | 0 | 0.00477 | CD38,CDKN1A,IGFBP3,PLAUR |
| CSF1 | 1.214 | 3.665 | 2.12E-08 | CCL2,CCR5,CD163,CD68,CDKN1A,CSF1,CSF1R,CTSK,FCER1G, IGF1,IL6,ITGAM,JUN,MMP16,MRC1,MSR1,PTPRC,TLR1 |
| CRH | 1.57 | 1.498 | 0.00185 | CRH,HPGD,IL1RN,IL6,STAR,UCN |

**Pi16-/-: Sham vs SNI**

| Upstream Regulator | Expr Log Ratio | Activation z-score | p-value of overlap | Target molecules in dataset |
|---|---|---|---|---|
| IL24 | 0.123 | _ | 0.0283 | GADD45A,GADD45G |
| TIMP1 | 0.231 | _ | 0.0158 | CDKN1A,PLAUR |
| CSF1 | 0.883 | 1.981 | 0.024 | CDKN1A,CSF1,MMP16,MRC1 |

**Figure S6, Related to Figure 7.** IPA's upstream regulator analysis was used to identify potential "cytokines" causing changes in gene expression in response to SNI in DRG between Sham and SNI group.

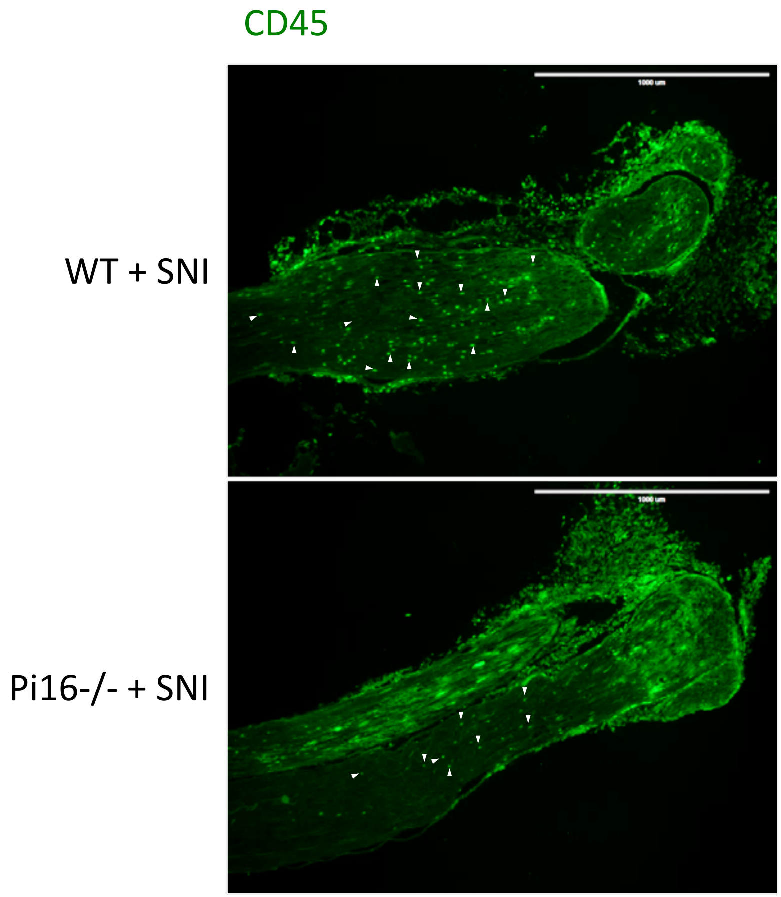


**Figure S7, Related to Figure 7.** Lower magnification image of sciatic nerve section with a larger field of view after SNI from the ipsilateral side (day 5) showing immunostaining of CD45 (green) in WT and Pi16-/- mice. White arrowheads indicate few of the CD45 positive cells. Note reduced number of CD45 positive cells (arrowheads) in Pi16-/- compared to WT. Scale bar indicates 1000 µm.
